# Chemical screens identify HDAC6 as an epigenetic vulnerability in acquired Temozolomide-resistant models of glioblastoma

**DOI:** 10.64898/2026.05.07.723296

**Authors:** Filiz Senbabaoglu Aksu, Buse Cevatemre, Nareg Degirmenci, Ezgi Yagmur Kala, Duygu Uçku, Martin Philpott, Adam P. Cribbs, James Dunford, Udo Oppermann, Ceyda Acilan, Tugba Bagci-Onder

## Abstract

Glioblastoma (GBM) is an aggressive primary brain tumor associated with a median survival of approximately 15 months following diagnosis. Current standard-of-care treatment includes surgical resection followed by radiotherapy and chemotherapy with the DNA-alkylating agent temozolomide (TMZ). However, tumor recurrence in a therapy-resistant state remains a major driver of poor patient outcomes. To investigate the molecular mechanisms underlying TMZ resistance, we generated *in vitro* models of acquired resistance by exposing initially TMZ-sensitive GBM cells to escalating doses of TMZ. Transcriptomic and chromatin accessibility profiling revealed extensive remodeling of DNA damage response (DDR) and DNA repair pathways that favored protection against TMZ-induced genotoxic stress. Although upregulation of O-6-methylguanine-DNA methyltransferase (MGMT) emerged as a dominant determinant of resistance in our models, the data suggested that additional adaptive resistance mechanisms contribute to the resistant phenotype. To identify targetable epigenetic dependencies associated with TMZ resistance, we performed a chemical screen using an epigenetic probe library. This screen identified multiple histone deacetylase (HDAC) inhibitors that selectively impaired the viability of TMZ-resistant cells, either as monotherapy or in combination with TMZ. Among these, HDAC6-selective inhibitors, including Ricolinostat, were particularly effective at inducing cell death in TMZ-resistant GBM models. Mechanistically, HDAC6 inhibition reduced the expression of key DDR-associated genes, while MGMT responses varied depending on cellular context and treatment duration. Furthermore, pharmacological inhibition and loss-of-function studies demonstrated that targeting HDAC6 could restore TMZ sensitivity by altering the balance of DNA repair pathway activity, potentially acting as a compensatory mechanism for MGMT-mediated resistance. Collectively, our findings identify HDAC6 as an epigenetic vulnerability in acquired TMZ-resistant GBM and support the therapeutic potential of HDAC6 inhibition as a strategy to overcome TMZ resistance in glioblastoma patients.

## INTRODUCTION

Glioblastoma (GBM) is the most common of all malignant brain and central nervous system tumors. Annually, the incidence of primary malignant brain tumors is about 7 per 100,000 individuals, increasing with age, and the five-year survival rate is approximately 36%, averaging across all types^1,2^. Current therapies for GBM include surgery, radiation therapy, and Temozolomide (TMZ) administration, with the combination of TMZ and radiotherapy resulting in a median survival of 14.6 months compared to 12 months for patients treated with radiotherapy alone^3^. This treatment option confers only a modest increase in patient survival and is limited by inherent or acquired therapeutic resistance of tumor cells^4,5^.

The development of TMZ resistance is a critical challenge in GBM treatment^6^. Well-established mechanisms of TMZ resistance include overexpression of O6-methylguanine-DNA methyltransferase (MGMT), which directly removes alkyl adducts from DNA, and deficiencies in the DNA mismatch repair (MMR) pathway^6^. However, TMZ resistance is a complex, multifactorial phenomenon that is not solely attributed to alterations in DNA repair mechanisms. Emerging evidence suggests that adaptive responses, including epigenetic rewiring, play a significant role in mediating resistance and promoting tumor survival^7^. Therefore, identifying novel targets and therapeutic strategies that can overcome or circumvent these complex resistance mechanisms is paramount for improving GBM patient survival.

Epigenetic changes play a key role in the pathobiology of GBM, encompassing disruptions in DNA methylation patterns and histone modification dynamics^8^. These alterations are driven by aberrant DNA methylation patterns, histone modifications, and changes in chromatin architecture^9^. Although several histone deacetylase inhibitors (HDACi) have shown promise in preclinical GBM models^10,11^, their clinical application remains limited. Ongoing trials primarily evaluate them in combination with TMZ, with class I HDAC inhibitor agents such as non-selective molecules like Vorinostat (NCT00268385) and Abexinostat (NCT05698524).

To identify epigenetic re-sensitizers that can overcome acquired TMZ resistance in GBM, we generated resistant cell line models and performed transcriptomic analyses, which revealed MGMT upregulation as a key contributor to the resistant phenotype. A focused screen of epigenetic compounds identified HDAC6 inhibitors as top hits selectively impairing the viability of resistant cells. HDAC6 inhibition disrupted DNA repair programs and restored TMZ sensitivity. CRISPR/Cas9-mediated knockout of HDAC6 phenocopied the effects of chemical inhibition, and similar sensitization was observed in primary patient-derived GBM cells, underscoring the therapeutic relevance of targeting HDAC6 in TMZ-resistant glioblastoma.

## MATERIALS AND METHODS

### Cell culture

Human GBM cell lines U373, T98G (CRL-1690), and A172 (CRL-1620) and HEK293T cells were purchased from ATCC. All cells except primary cells were cultured in DMEM medium (Gibco, USA) with 10% fetal bovine serum (Invitrogen, USA) and 1% Penicillin-Streptomycin (Gibco, USA). The protocol for establishing primary cell lines from GBM patient biopsies has been previously described^12^. These cells were grown in Neurobasal medium (Gibco, 21103-049) containing L-Glutamine (Gibco, 25-030-081), B27 and N2 supplements (Gibco, 17-504-044, A1370701), 0.5% Penicillin-Streptomycin (Gibco, 15-070-063), Heparin (Stem Cell Technologies, 07980), FGF (20 ng/ml, Gibco, #PHG0266), and EGF (20 ng/ml, Peprotech, #AF-100-15). Cells were maintained in a humidified incubator at 37°C with 5% CO2. Routine checks for mycoplasma contamination were performed using the Lonza Mycoplasma Detection Kit (#LT07-318).

### Reagents

The drug library, consisting of 96 epigenetic probes targeting various histone-modifying enzymes (Table S1), was described previously^13^; it comprises chemical probes from the Structural Genomics Consortium together with reference inhibitors against targets not covered by the SGC probe set (including HDAC inhibitors). Temozolomide (S1237), CAY10603 (S7596) and Lomustine (S1840) were purchased from SelleckChem. Ricolinostat (21531), Cisplatin (21531) and Carboplatin (13112) were obtained from Cayman Chemicals. The pJP1520 MGMT overexpression plasmid was purchased from DNAsu (Clone: HsCD00075543) and transduced into U373 cells.

### Establishment of TMZ-resistant cells

The TMZ-resistant U373 cells were established using methodology adapted from Banelli et al.^12^, the dose escalation method. U373 cells were treated with TMZ every 3 days, starting at 25 μM. The dosage was doubled every two weeks (or when the cells became resistant to the current dose), and these cells were denoted as DE-TMZR. In parallel, a second resistant population (TMZR) was established by a single-pulse regimen (250 µM TMZ) for 15 days followed by 15 days of drug-free recovery. A derivative of this cell line which is cultured under TMZ maintenance dose is described previously^14^. U373 parental cells were maintained as age-matched controls alongside the TMZ-resistant cells for comparison.

### Cell viability assays

Cells were seeded at 1000 cells per well in black 96-well plates and treated with the respective drug the following day for 3 or 5 days. Cell viability was measured using the ATP-based Cell Titer-Glo (CTG) Luminescent Cell Viability Assay (Promega) according to the manufacturer’s instructions. Relative luminescence unit (RLU) values were obtained using a plate reader (Synergy H1 Reader, Bio-Tek). For long-term viability evaluation, a clonogenic assay was performed. Briefly, cells were seeded in triplicate at 0.5– 1 × 10^3^ cells per well in 6-well plates. The next day, the drug was administered at varying doses for 48 h. After 10–14 days, cells were fixed with cold methanol and stained with crystal violet. Colony quantifications were performed using ImageJ.

### γH2AX activation for DNA damage

H2A.X activation was measured using the Muse™ H2A.X Activation Dual Detection Kit (Millipore, #MCH200101), following the manufacturer’s instructions. Cells were treated with 250 µM of DMSO or TMZ for 24 h, and phospho-H2AX levels were detected using the Muse™ Cell Analyzer.

### Host Cell Reactivation (HCR) assay

The pBOS-H2BGFP DNA (1.5 µg per treatment group) was incubated with TMZ at various concentrations (0–5 mM) in TE buffer at room temperature for 24 h. The TMZ-damaged plasmid was purified by NucleoSpin PCR clean-up kit according to manufacturer’s instructions (Macherey-Nagel, 740609). Cells were seeded at 200,000 per well in 6-well plates and transfected using Lipofectamine 3000 (1.5 µg DNA/3.75 µL Lipofectamine, Life Technologies, L3000015) according to the manufacturer’s instructions. At 24, 48, and 72 h post-transfection, cells were dissociated with trypsin, washed with PBS, and approximately 1 × 10^4^ cells per sample were acquired. GFP expression levels were measured using the BD Accuri C6 flow cytometer (BD Bioscience), and DNA repair capacity was calculated as a percentage of the GFP signal from cells transfected with undamaged plasmid.

### Epigenetic drug screen

Cells were seeded at 1,250 per well in black 96-well plates. The following day, the cells were treated with the chemical inhibitors at their respective working concentrations (determined by SGC, Table S1) and/or TMZ (0.25 mM for U373, and 0.5 mM for DE-TMZR). These conditions were not equitoxic across models and served as an initial screening framework. Cell viability was detected after 72 h using the CTG assay, and the results were normalized to untreated controls.

### RNA-sequencing

RNA was isolated from cells treated with (36 h) or without Ricolinostat using the Quick-RNA Mini prep kit (Zymo Research, R1054) following the manufacturer’s protocol. The quality of the RNA samples was verified by electrophoresis on TapeStation (Agilent). The RNA integrity number (RIN) scores for all samples ranged from 7.5 to 9.5. RNA-seq libraries were prepared using the NEBNext® Ultra™ RNA library prep kit for Illumina® with TruSeq indexes, according to the manufacturer’s protocol. The resulting libraries were sequenced on a NextSeq 500 platform (Illumina) using a paired-end run of 2 × 80 bp, achieving an average depth of 112 × 10^6^ paired-end reads per sample (ranging from 47 × 10^6^ to 168 × 10^6^). Analysis was performed on Genialis Expressions Platform (Genialis, Inc., Boston, MA, USA).

### ATAC-sequencing

ATAC-seq was performed using 100,000 cells for the transposition reaction, which was performed based on the protocol by Buenrostro *et al*.^15,16^ using in-house produced transposase. Briefly, cells were washed with cold PBS, lysed in 100 µl cold lysis buffer containing 10 mM Tris-HCl pH 8, 10 mM NaCl and 4 mM MgCl2, 0.3% Nonidet NP40 and 0.01% Tween 20 for 5 min. After centrifugation of 10 min 500 g at 4°C, cells were resuspended in TD Buffer containing 0.1% Tween 20 and 0.05% Digitonin for 5 min at room temperature. After addition of 4 μl Tn5 enzyme, incubation at 37°C for 60 min with 500 rpm shaking was performed. Subsequently, the samples were purified. PCR-amplification was performed using the following protocol: 3 min 72°C; 30 sec 98°C; 11 cycles of (10 sec 98°C, 30 sec 63°C, 3 min 72°C). Primers are listed in Table S3. The samples were then purified and eluted with 20 μL of TE buffer. Samples were then validated on a TapeStation (Agilent) to determine library size and quantification prior to paired-end (2 × 41 bp) sequencing on a NextSeq 500 (Illumina) platform. Adapter sequences used in ATAC-seq are listed in **Table S2**.

### Quantitative real-time PCR (RT-qPCR)

RNA was collected using an RNA isolation kit following the manufacturer’s instructions (Macherey-Nagel, 740955). A total of 1 μg RNA was used to synthesize cDNA with M-MLV Reverse Transcriptase (Invitrogen, 28025013). Relative gene expression levels were detected using LightCycler 480 SYBR Green I Master (Roche, 04707516001). The primers are provided in **Table S3**.

### CRISPR/Cas9

To establish CRISPR-Cas9-mediated knockout cells, we followed the protocol of Zhang Lab^16^ and cloned gRNAs targeting MGMT or HDAC6 into the pLenti-CRISPRv2 plasmid. The oligonucleotides are provided in **Table S3**. Validation of the knockout was carried out using WB analysis.

### Western blotting (WB)

WB analysis was performed as previously described^17^. Briefly, for whole cell extracts, a buffer was prepared containing 1% NP-40, 150 mM NaCl, 1 mM EDTA, 50 mM Tris-HCl (pH 7.8), 1 mM NaF, 0.5 mM PMSF, 1× phosphatase inhibitor cocktail (PhosSTOP, Roche, #11699695001), and 1× protease inhibitor cocktail (cOmplete Protease Inhibitor Cocktail Tablets, Roche, #11697498001). Protein concentration was measured using the Pierce BCA Protein Assay Kit (Thermo Fisher, #23225) following the manufacturer’s instructions. Equal amounts of protein were loaded onto polyacrylamide gels, and proteins were transferred onto the Immun-Blot PVDF membrane (Bio-Rad) by the Trans-Blot Turbo Transfer System (Bio-Rad). The antibodies used and their respective dilutions are listed in **Table S4**.

### Patient data analysis

Vital status data were downloaded via cBioPortal^18^ using the Glioblastoma^19^ (CPTAC, Cell 2021) dataset, scatter-plotted in GraphPad and analyzed with a t-test. The response of astrocytoma patients receiving TMZ treatment was accessed through CTR-DB^20^; plots and additional details (sample size, logFC, logFC p-value) were obtained directly from the site. The correlation between HDAC6 and the ATM and ATR genes was analyzed using the Glioblastoma^21^ (TCGA, Cell 2013) dataset, downloaded via cBioPortal, and the correlation analysis was performed using GraphPad.

### Statistical analysis

All normalizations were performed using GraphPad Prism version 8.0, and statistical analyses were specified in the figure legends.

## RESULTS

### Generation and characterization of TMZ-resistant cell lines

We derived resistant cell populations from the TMZ-sensitive U373 cell line using the dose-escalation approach, where we started with 25 µM and doubled the dose up to 250 µM over three months (**Fig. 1A**). These TMZ-resistant cells, named “DE-TMZR”, were confirmed to be resistant up to 500 µM using clonogenic assays (**Fig. 1B**). In parallel, we established a second resistant population (TMZR) by a single-pulse regimen, 250 µM TMZ for 15 days followed by 15 days of drug-free recovery and confirmed resistance by clonogenic survival across 125–500 µM TMZ (**Fig. S1A, B**). As expected, the TMZ-resistant cells showed higher IC50 values compared to the parental cells (71 µM vs. 953 µM and 1532 µM respectively, **Fig. 1C, Fig. S1C**). Additionally, a slower proliferation rate was observed in the resistant cells (**Fig. 1D**).

**Figure 1.**
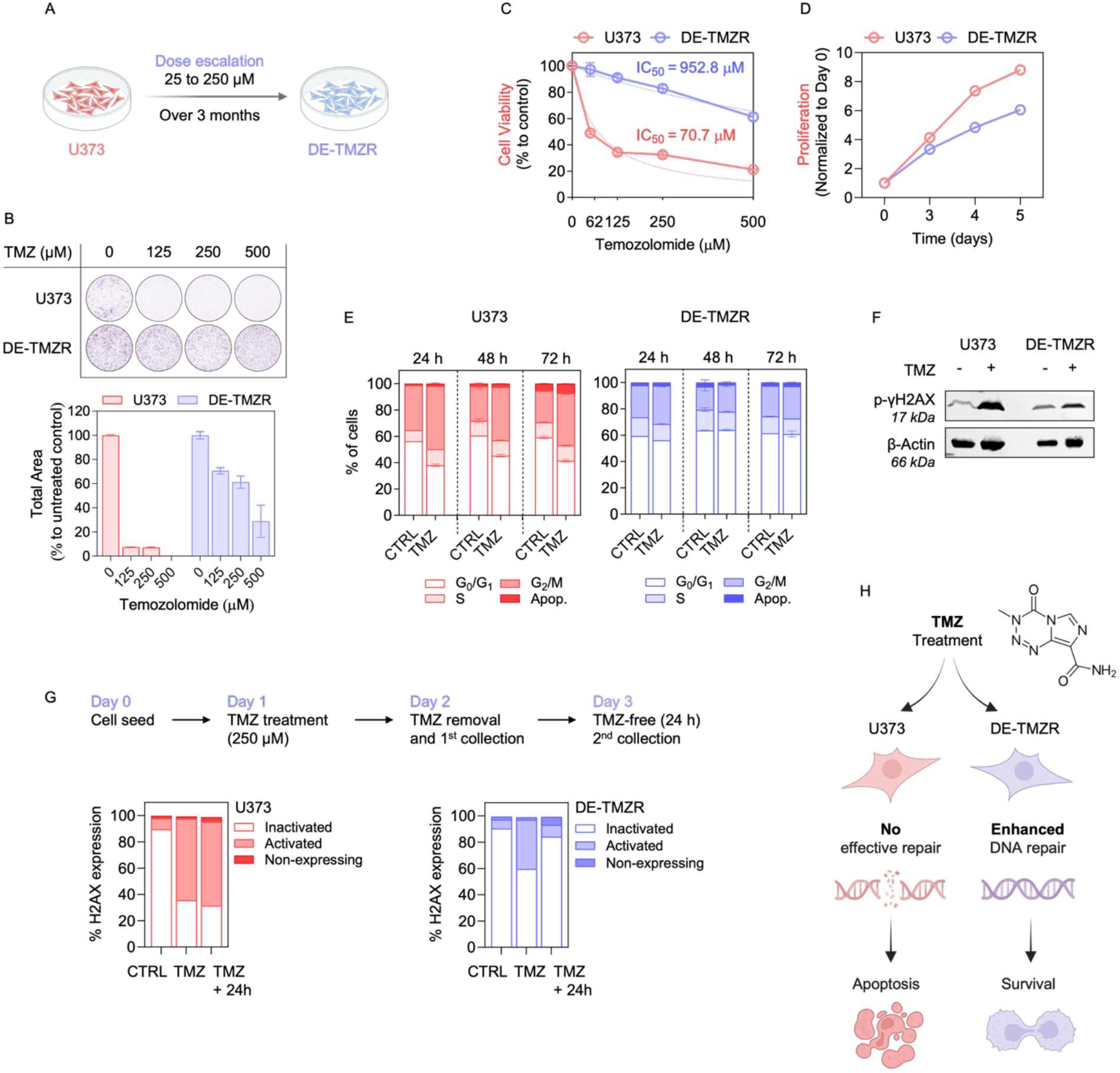
Establishment and characterization of TMZ-resistant U373 cell lines. **A**. Schematics of the establishment of TMZ-resistant cells (DE-TMZR) using the dose escalation method. Cells were treated with TMZ every 2-3 days, with the concentration doubled every 2 weeks. Created in BioRender. **B**. Cells were treated with TMZ for 48 h, and after drug removal, they were allowed to proliferate for 10 days. Colony areas were measured and normalized to their respective controls. **C**. Cell Titer Glo (CTG) cell viability results were obtained after 5 days of TMZ treatment. **E**. Proliferation curves of U373 and DE-TMZR cells. Cell cycle analysis was performed by flow cytometry using propidium iodide staining after 24 h of TMZ treatment (250 µM). All assays were performed in triplicate, and error bars represent the standard deviation. **F**. γ-H2AX protein levels were assessed by Western blot analysis after TMZ treatment (250 µM, 24 h); β-actin is used as a loading control. **G**. The γ-H2AX level was determined using the Muse flow cytometer. **H**. BioRender illustration showing the proposed mechanism of acquired TMZ resistance in DE-TMZR cells. Upon TMZ treatment, DE-TMZR cells repair DNA damage and survive, whereas parental U373 cells fail to repair and undergo apoptosis.

TMZ exerts its cytotoxic effects primarily through DNA damage, particularly by inducing double-strand breaks (DSBs), which in turn activate DNA damage response pathways and trigger cell cycle arrest^22^. To investigate how this mechanism manifests in our models, we first analyzed TMZ-induced changes in the cell cycle using flow cytometry. Under identical dosing, U373 underwent G2/M arrest while DE-TMZR did not (**Fig. 1E**). To assess the extent of DNA damage induced by TMZ, γ-H2AX levels were analyzed by Western blotting. While γ-H2AX induction was pronounced in parental U373 cells, the signal was markedly lower in both DE-TMZR and TMZR cells (**Fig. 1F, Fig. S1D**). To address the capacity to repair DSBs, the activation of the γ-H2AX protein was examined. Cells were treated with 250 µM TMZ for 24 h, and γ-H2AX levels were measured immediately after treatment and 24 h following drug removal (**Fig. 1G**). TMZ treatment induced a significant increase in γ-H2AX levels in the parental cell line compared to DE-TMZR cells, and these elevated γ-H2AX levels remained unchanged after drug removal. In DE-TMZR cells, we observed a partial increase in activated γ-H2AX following TMZ treatment, which was reversed after drug removal (unlike in parental cells), indicating differences in the repair capacities of the cells (**Fig. 1H**). To further assess DNA repair capacity in U373 and DE-TMZR cells, we performed the Host Cell Reactivation (HCR) assay. This transfection-based assay evaluates the ability of host cells to repair damage to exogenous DNA (**Fig. S2A**). In our study, the HCR assay involved the use of TMZ-damaged or undamaged plasmids encoding the GFP gene. Since TMZ-induced DNA lesions block transcription, these lesions must be repaired for GFP expression to occur. Thus, the GFP signal indicates the extent of DNA repair following transfection and recovery of the plasmid in the cells. We observed the enhanced DNA repair capacity in DE-TMZR cells (**Fig. S2B**). To test whether our established TMZ-resistant cell lines were also cross-resistant to other DNA alkylating agents, we treated the cells with Cisplatin, Carboplatin, and Lomustine; Cyclophosphamide was profiled without metabolic activation (**Fig. S3**). No significant differences were observed for Cisplatin and Carboplatin; Cyclophosphamide requires metabolic activation and was not interpretable under these conditions. We noted a comparable resistance phenotype for Lomustine in our TMZ-resistant cell lines (**Fig. S3**), which was consistent with the similar mechanisms of action shared by Lomustine and TMZ.

### Transcriptome analysis reveals MGMT as one of the top upregulated genes in TMZ-resistant cell lines

Transcriptional programs can be rewired during the adaptation of cells to drug treatment and the acquisition of drug resistance. To investigate these changes, we conducted RNA- and ATAC-sequencing on resistant cells (vs. parental) to uncover key transcriptional and epigenetic alterations. Integration of the two datasets revealed 318 overlapping genes (**Fig. 2A**, left). From this intersection, we ranked the genes by their expression changes, and *MGMT* emerged among the top upregulated genes (**Fig. 2A**, right). *MGMT* is also upregulated in TMZR and ranks within the top ten upregulated genes shared with DE-TMZR (**Fig. S4A, B**). ATAC-seq profiling indicated an open chromatin configuration at the *MGMT* locus in DE-TMZR cells, particularly around the transcription start site (TSS; chromosome 10:131265454–131565783, RefSeq NM_002412), consistent with the transcriptional activation observed in RNA-seq (**Fig. 2B**). Validation by RT-qPCR and Western blot confirmed robust MGMT expression in DE-TMZR cells, whereas parental U373 cells exhibited undetectable levels (**Fig. 2C**).

**Figure 2.**
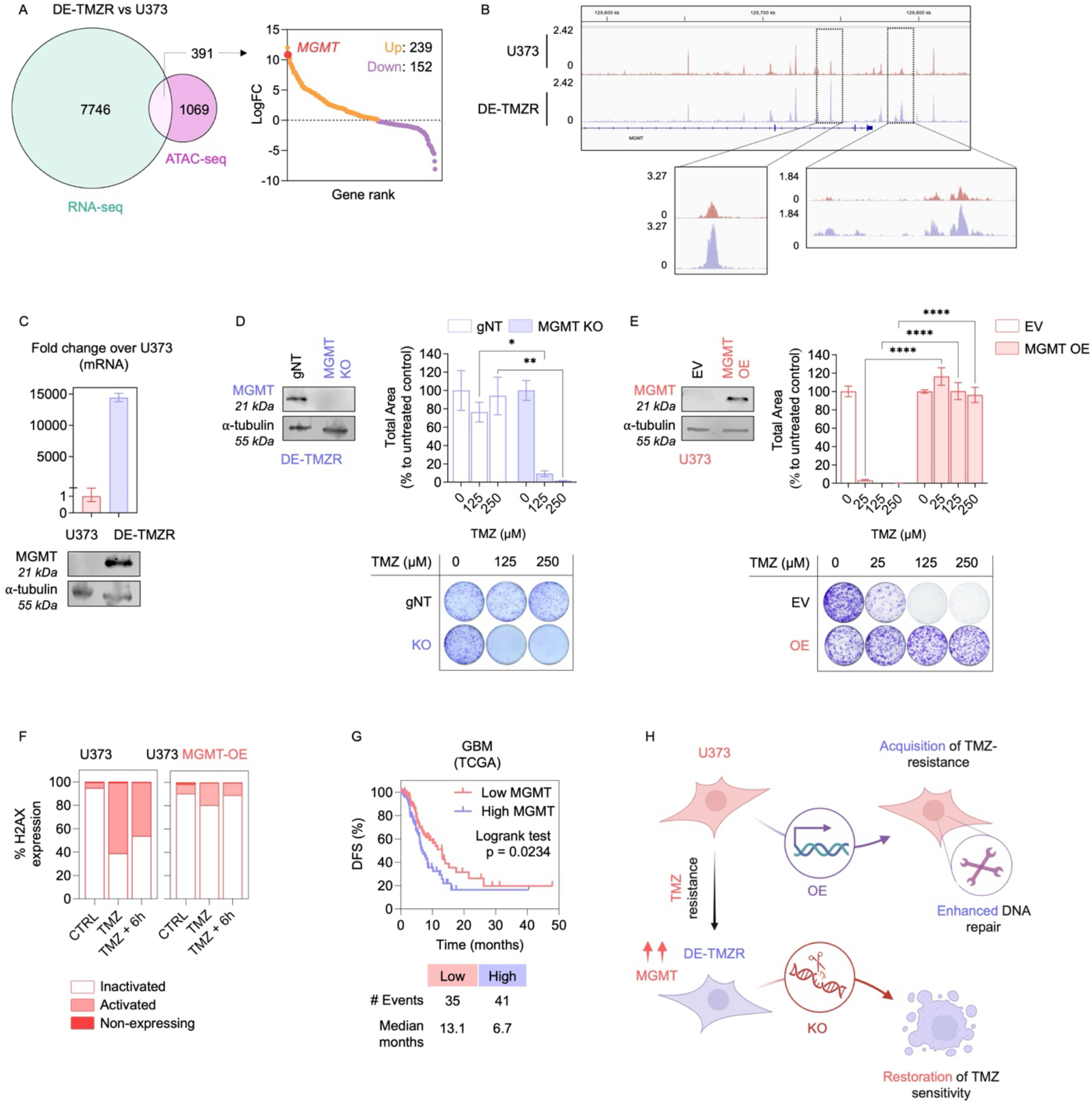
Integrated transcriptomic and chromatin accessibility analyses reveal MGMT dependency in TMZ response. **A**. Venn diagram showing the overlap between differentially expressed genes (RNA-seq) and altered chromatin-accessible regions (ATAC-seq) in DE-TMZR versus U373 cells. Among the 391 common genes, *MGMT* was highlighted as one of the top upregulated genes. **B**. Representative ATAC-seq peak signal illustrating enhanced chromatin accessibility at the *MGMT* locus in DE-TMZR cells relative to U373. **C**. Validation of increased *MGMT* expression in DE-TMZR compared with U373 cells by RT-qPCR (up) and immunoblotting (down). **D**. MGMT protein levels and clonogenic assay results of DE-TMZR cells following MGMT knockout, with corresponding quantification of colony formation efficiency. **E**. MGMT protein levels and clonogenic assay results of U373 cells following MGMT overexpression (OE), with corresponding quantification of colony formation efficiency. **F**. DNA damage was assessed by quantifying phospho-H2AX (γH2AX) expression in U373 and MGMT-overexpressing (MGMT-OE) cells following 250 µM TMZ treatment. Cells were analyzed 6 h after drug removal using Muse flow cytometry. **G**. Kaplan-Meier analysis of disease-free survival (DFS) in GBM patients from the TCGA dataset stratified by *MGMT* expression levels. Patients with high MGMT expression exhibit significantly reduced DFS compared to those with low MGMT expression (log-rank p = 0.0234). **H**. BioRender schematic depicting the acquisition and restoration of TMZ sensitivity. TMZ exposure induces resistance in U373 cells (DE-TMZR) through MGMT upregulation, leading to enhanced DNA repair and increased TMZ tolerance. MGMT knockout reverses this phenotype and restores TMZ sensitivity, resulting in cell death.

To determine the functional relevance of MGMT induction, we used a loss-of-function approach to knock out MGMT in both DE-TMZR and TMZR cells. MGMT-KO cells displayed a marked decrease in clonogenic survival following TMZ exposure compared with controls (**Fig. 2D, Fig. S4C**). Conversely, MGMT-overexpressing U373 cells exhibited resistance to TMZ, comparable to the resistance levels observed in our TMZ-resistant cells (**Fig. 2E**). Additionally, when MGMT-overexpressing U373 cells were treated with TMZ for 24 h, we observed a markedly attenuated induction of phospho-H2AX compared to the parental U373 cells, consistent with a reduced DNA damage response upon TMZ exposure (**Fig. 2F**). Following drug removal, phospho-H2AX levels rapidly returned to baseline in MGMT-overexpressing cells (**Fig. 2F**). These findings support MGMT as a dominant contributor to TMZ resistance in our U373-derived TMZ-resistant models.

To further evaluate the clinical relevance of MGMT expression, analysis of the TCGA-GBM cohort revealed that patients with high MGMT levels exhibited significantly shorter disease-free survival compared to those with low MGMT expression (median = 6.7 vs. 13.1 months; p = 0.0234, **Fig. 2G**). Consistently, in the CPTAC dataset, MGMT mRNA levels were significantly higher in deceased GBM patients relative to living cases (p = 0.0189, **Fig. S5A**). Similarly, analysis of CTR-DB data showed that astrocytoma patients receiving TMZ treatment with high MGMT expression were predominantly non-responders (**Fig. S5B**). The evidence supports that MGMT plays a central part in enabling DNA repair and promoting TMZ resistance in GBM cells, as illustrated in the schematic model (Fig. 2H), and further demonstrate that its overexpression not only protects against TMZ-induced DNA damage but also correlates with poorer clinical outcomes.

### Chemical screen identifies HDAC6 inhibitor Ricolinostat as an effective epigenetic drug in TMZ-resistant GBM cells

To identify epigenetic compounds with therapeutic potential in TMZ-resistant GBM, we performed a compound screen using an epigenetic probe library that targeted chromatin modifiers such as writers, erasers and reader proteins (**Fig. 3A**) with a total of 97 compounds. Cells were treated either with the library compounds alone or in combination with TMZ, with the threshold for hits set at 50% cell viability when combined with TMZ (**Fig. 3B**). Notably, two independent PARP inhibitors, Olaparib and Rucaparib, scored among the top hits (**Fig. 3C**) and enhanced TMZ activity in both parental and resistant lines, with more pronounced effects in the resistant derivatives (**Fig. S6A–D**), consistent with a proof-of-principle demonstration.

**Figure 3.**
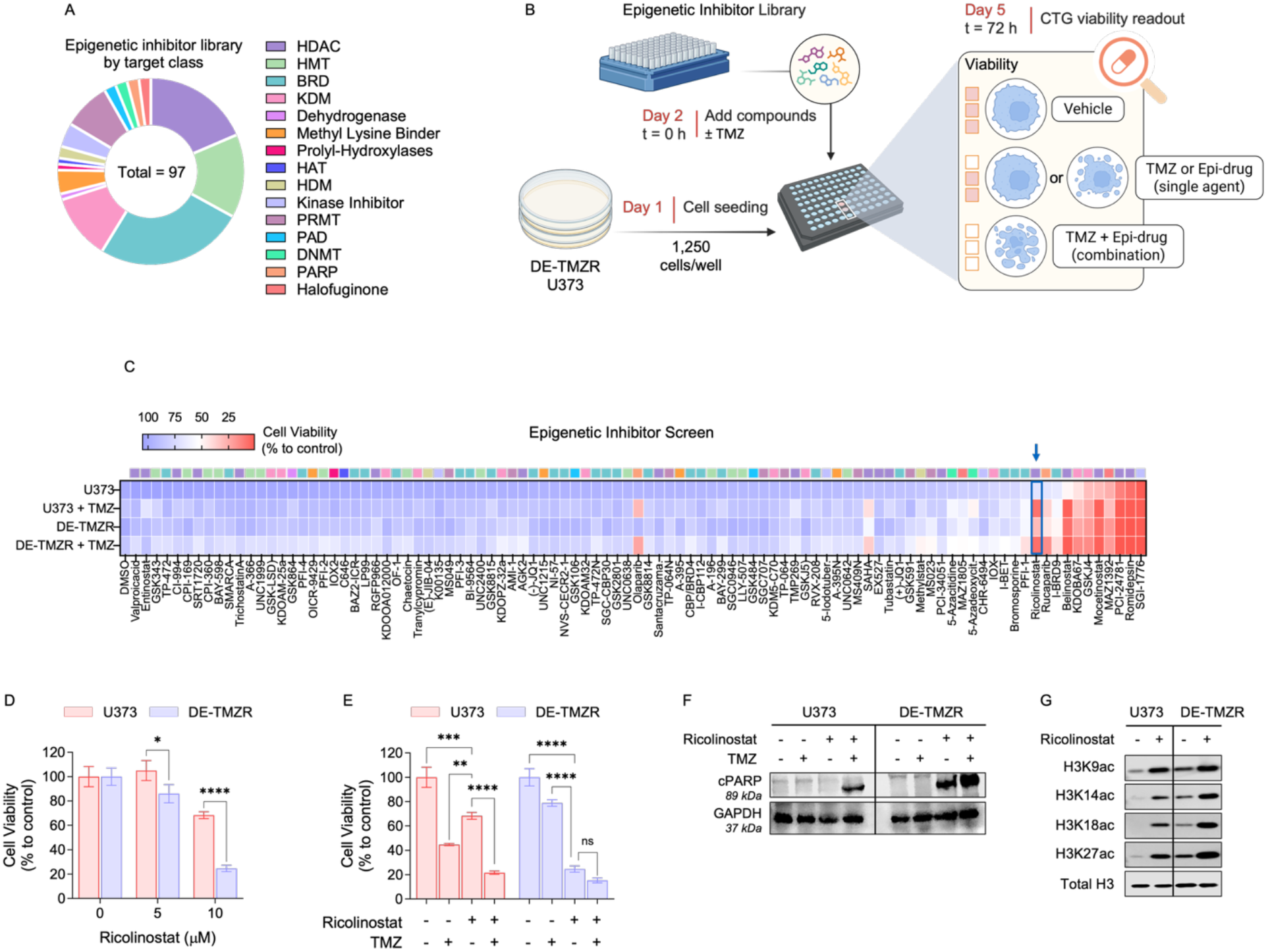
A drug screen of epigenetic inhibitors reveals HDAC6 inhibitor Ricolinostat as a TMZ sensitizer. **A**. Composition of the epigenetic inhibitor library (total = 97 compounds) grouped by target class. **B**. Screening workflow created with BioRender. U373 and DE-TMZR cells were seeded at 1,250 cells/well (Day 1) and, on Day 2 (t = 0 h), treated with each epigenetic compound alone or with TMZ, alongside vehicle controls. Viability was measured on Day 5 (72 h) using CTG. **C**. Cell viability results from screens performed on U373 and DE-TMZR cells. Cells were treated with the respective drug, with or without TMZ (250 µM for U373 and 500 µM for DE-TMZR) for 72 h. Cell viability was measured using the CTG assay and normalized to untreated controls. **D**. Dose-dependent viability results of Ricolinostat treatment for 72 h. **E**. Cell viability results of the combination of Ricolinostat (10 µM) and TMZ (250 µM). **F**. Western blot showing PARP cleavage following 36 h treatment with Ricolinostat (10 µM) and/or TMZ (250 µM), with GAPDH used as a loading control. **G**. Western blot analysis of Histone 3 acetylation marks after 36 h of Ricolinostat (10 µM) treatment, with total H3 as a loading control.

To determine whether the sensitization observed in TMZ-resistant GBM cells is a generalizable phenomenon, we additionally evaluated T98G glioblastoma cells, which exhibit innate TMZ resistance driven by high MGMT expression (**Fig. S7A**). T98G cells displayed minimal reduction in clonogenic survival upon TMZ treatment, and MGMT-KO markedly increased TMZ sensitivity, confirming the functional contribution of MGMT in intrinsic resistance (**Fig. S7B**). We next performed a focused epigenetic inhibitor screen in T98G cells using the same library. In contrast to the sensitization observed in U373 and DE-TMZR, compound treatment alone or in combination with TMZ failed to induce substantial loss of viability in T98G cells (**Fig. S7C**). These results indicate that intrinsic MGMT-driven resistance in T98G cells does not confer susceptibility to epigenetic perturbation, unlike the epigenetically rewired acquired-resistant models generated in this study.

Among the hits, the majority clustered as HDAC inhibitors (**Fig. 3C**). In general, the acetylation of histone lysines promotes an open chromatin structure, which is linked to active gene transcription, while histone deacetylation leads to a more compact chromatin state, indicating transcriptional repression^23^. Thus, the balance between HATs and HDACs is crucial for regulating gene expression. Elevated HDAC expression can promote tumor development^24^. In GBM, HDACi have shown promising preclinical potential, suggesting that HDACi treatment sensitizes tumor cells to chemotherapy and induces apoptosis^25,26^. Pan-HDAC inhibitors such as Vorinostat (NCT03426891) and Quisinostat (NCT06824662) are currently undergoing clinical trials for GBM patients.

Among HDACi hits, Ricolinostat (ACY-1215), a selective HDAC6 inhibitor, was highlighted in **Fig. 3C** as it showed stronger effects in TMZ-resistant cells (viability < 50%) compared to the parental U373 cells under the tested conditions (**Fig. 3D, Fig. S8A**). We further validated the screening results for Belinostat and another HDAC6 inhibitor, CAY10603, confirming dose-dependent reduction of cell viability and the greater sensitivity of TMZ-resistant cells to HDAC6 inhibition compared with the parental counterpart (**Fig. 3E, Fig. S8A, Fig. S9A**). Consistent with these findings, apoptosis analysis revealed marked cPARP induction in Ricolinostat-treated TMZ-resistant cells but not in parental U373 cells, while the TMZ + Ricolinostat combination further enhanced cPARP levels, indicating restored chemosensitivity and apoptosis induction in resistant cells (**Fig. 3F, Fig. S8B**). Ricolinostat treatment led to a pronounced increase in H3K9ac, H3K14ac, H3K18ac, and H3K27ac levels, consistent with effective HDAC inhibition and confirming on-target activity (**Fig. 3G, Fig. S8C**).

### Ricolinostat downregulates DNA-repair gene expression and enforces cell-cycle checkpoint activation in parental and TMZ-resistant GBM cells

To examine mechanisms underlying the Ricolinostat response in GBM, we profiled parental U373 and TMZ-resistant derivatives (TMZR, DE-TMZR). Differential expression analysis (FDR < 0.05) revealed shared downregulated genes across all three lines (3,843 genes; **Fig. 4A**). Enrichment mapping highlighted clusters related to cell-cycle and DNA-repair/replication programs (**Fig. 4B**), and a chord diagram showed broad suppression of repair terms, with *ATM* appearing in multiple categories (**Fig. 4C**). Concordantly, ATM protein decreased after 36 h of Ricolinostat in U373, TMZR, and DE-TMZR, thereby validating the transcriptomic downregulation at the protein level (**Fig. 4D**). RNA-seq at a single 36 h time point and fold-change-based enrichments were initially reported without FDR; late apoptotic effects may contribute to pathway suppression. In addition to the ATM decrease, we confirmed RNA-seq-identified changes at the mRNA level by RT-qPCR (**Fig. S10A**). Consistently, Hallmark GSEA across the three lines showed common downregulation of cell-cycle and DNA-repair pathways (**Fig. S10B**). As a pharmacologic point of reference for ATM modulation, dose response cell-viability assays with the inhibitors KU55933 (ATM) and AZD6738 (ATR) resulted in more pronounced loss of viability in DE-TMZR/TMZR than in U373 (**Fig. S9B**), suggesting that the resistant derivatives may be more dependent on ATM/ATR signaling, in line with their elevated DNA-repair capacity (**Fig. 1G, Fig. S2B**).

**Figure 4.**
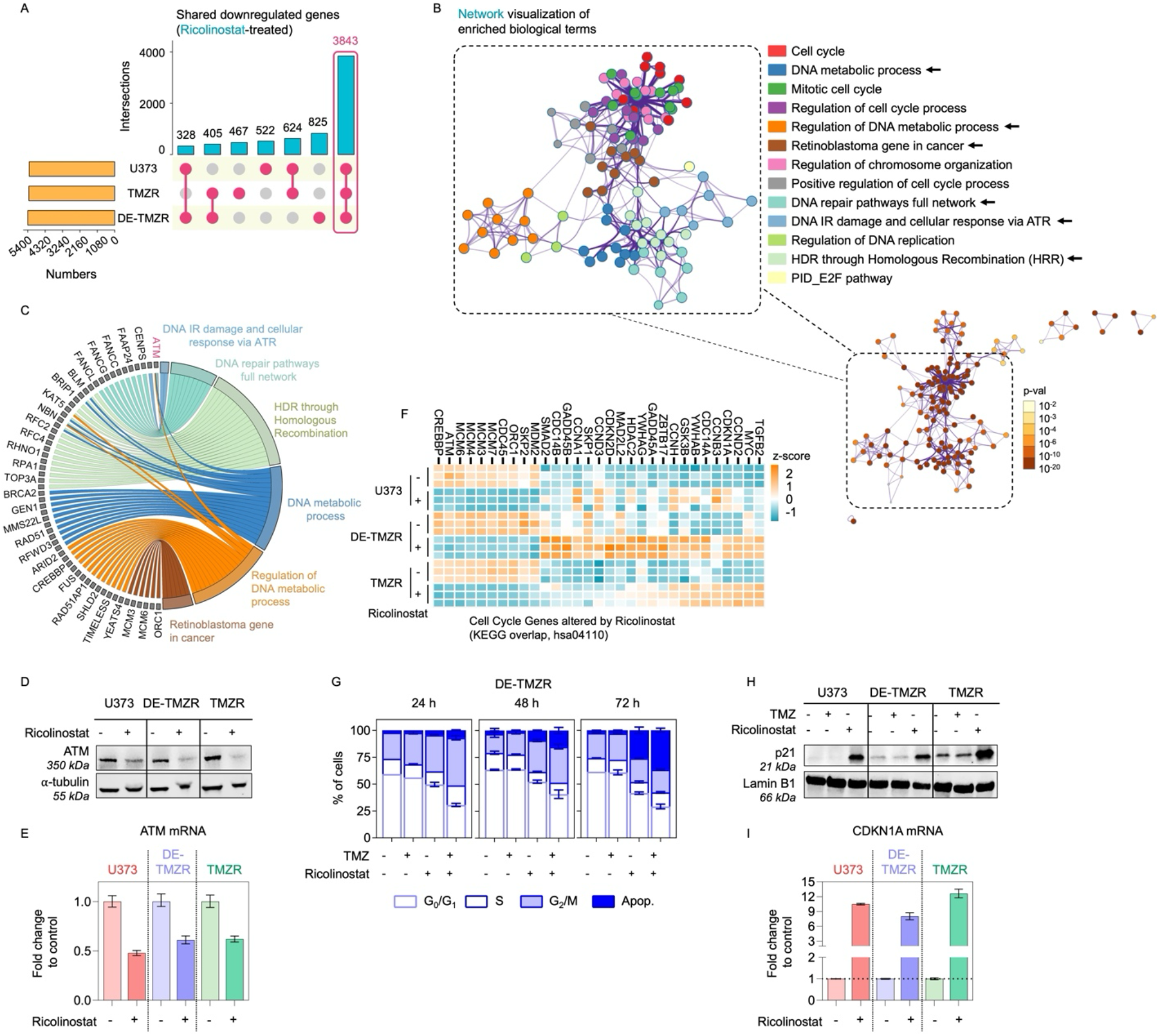
Transcriptomic profiling following Ricolinostat treatment reveals suppression of cell-cycle and DNA-repair programs. **A**. Shared downregulated genes across three Ricolinostat-treated cell lines (U373, TMZR, and DE-TMZR) visualized by an UpSet plot. **B**. Network of enriched biological terms obtained from Metascape analysis of the overlapping genes shown in (A) (n = 3843). The top 3000 genes, ranked by fold change (based on the DE-TMZR dataset), were subjected to enrichment analysis. Node size and color indicate p-value significance, and the zoom-in highlights the most significant subnetworks, colored by cluster ID. **C**. Chord diagram linking the shared downregulated genes to DNA-repair-related terms; colors match panel B. Ribbons denote gene-term membership. ATM appears across multiple DNA-repair categories, highlighted. **D**. Immunoblot validation of ATM downregulation following Ricolinostat treatment (36 h) in U373, TMZR, and DE-TMZR. α-tubulin is used as a loading control. **E**. RT-qPCR validation of *ATM* found to change 36 h after Ricolinostat across U373, DE-TMZR, and TMZR **F**. Heat map representing differentially expressed cell cycle genes with Ricolinostat treatment in U373, TMZR and DE-TMZR. **G**. Time-course cell-cycle profiling after Ricolinostat (10 µM) ± TMZ (250 µM) in DE-TMZR cells. **H**. Immunoblot showing induction of p21 after Ricolinostat treatment (24 h), consistent with cell-cycle arrest. Lamin B1 is used as a loading control. **I**. RT-qPCR validation of *CDKN1A* found to change 24 h after Ricolinostat across U373, DE-TMZR, and TMZR.

Among the downregulated enrichments, cell-cycle programs emerged as the other dominant theme in the network map (**Fig. 4B**). Consistent with this, intersecting DEGs with the KEGG Cell Cycle set revealed Ricolinostat-responsive cell-cycle genes shared across all three lines (**Fig. 4E–F**). Functionally, (±TMZ) profiling revealed G2/M accumulation at 24 h and subsequent apoptosis at later time points; co-treatment further enhanced these effects, consistent with Ricolinostat-mediated TMZ sensitization (**Fig. 4G**; see also **Fig. S10C–D** for U373 and TMZR). To focus on regulators, we intersected DEGs with the KEGG Cell Cycle set and plotted Ricolinostat-upregulated regulators shared across all three lines; among these, *CDKN1A* (p21) was robustly induced, which we validated by immunoblotting (**Fig. 4H–I**). Together, these data indicate that Ricolinostat suppresses DNA-repair programs and is consistent with checkpoint engagement; enforces cell cycle checkpoint activation in both parental and TMZ-resistant GBM cells.

### HDAC6 depletion enhances TMZ sensitivity in resistant and primary GBM

We performed a targeted siRNA screen against HDAC1/2/3/6/8 to test whether genetic depletion could phenocopy drug-mediated resensitization. Knockdown efficiency was confirmed for each target using two independent siRNAs per gene, showing efficient depletion relative to siControl (**Fig. S11A**). HDAC6 knockdown recapitulated the pharmacologic phenotype observed with HDAC6 inhibitors (Ricolinostat, CAY10603, Belinostat, **Fig. S8A, Fig. S11B**), and given the transient nature of siRNA, we next established stable cells with HDAC6 knockout using CRISPR/Cas9 (**Fig. 5A**). There was a slight, albeit significant, increase in TMZ sensitivity in U373 and DE-TMZR cells (**Fig. 5B**). To address these effects in a clinically-relevant model, we validated our findings in the patient-derived primary GBM cell line we previously established^27^ (**Fig. 5C**). We observed that HDAC6 inhibition resulted in a similar phenotype in KUGBM8-EF cells; Ricolinostat treatment increased TMZ responses, with an increase in *CDKN1A* expression and a reduction in *MGMT* expression (**Fig. 5D, E**). The knockout of HDAC6 similarly increased the H3K27ac levels and TMZ response. Genetic ablation of HDAC also led to an increase in *CDKN1A* expression and decrease in *MGMT* expression (**Fig. 5G, H**). Building on the results, the analysis of TCGA Glioblastoma datasets shows significant correlation between *HDAC6* and *ATM/ATR* expression, suggesting that HDAC6 may interface with ATM/ATR-mediated repair pathways.

**Figure 5.**
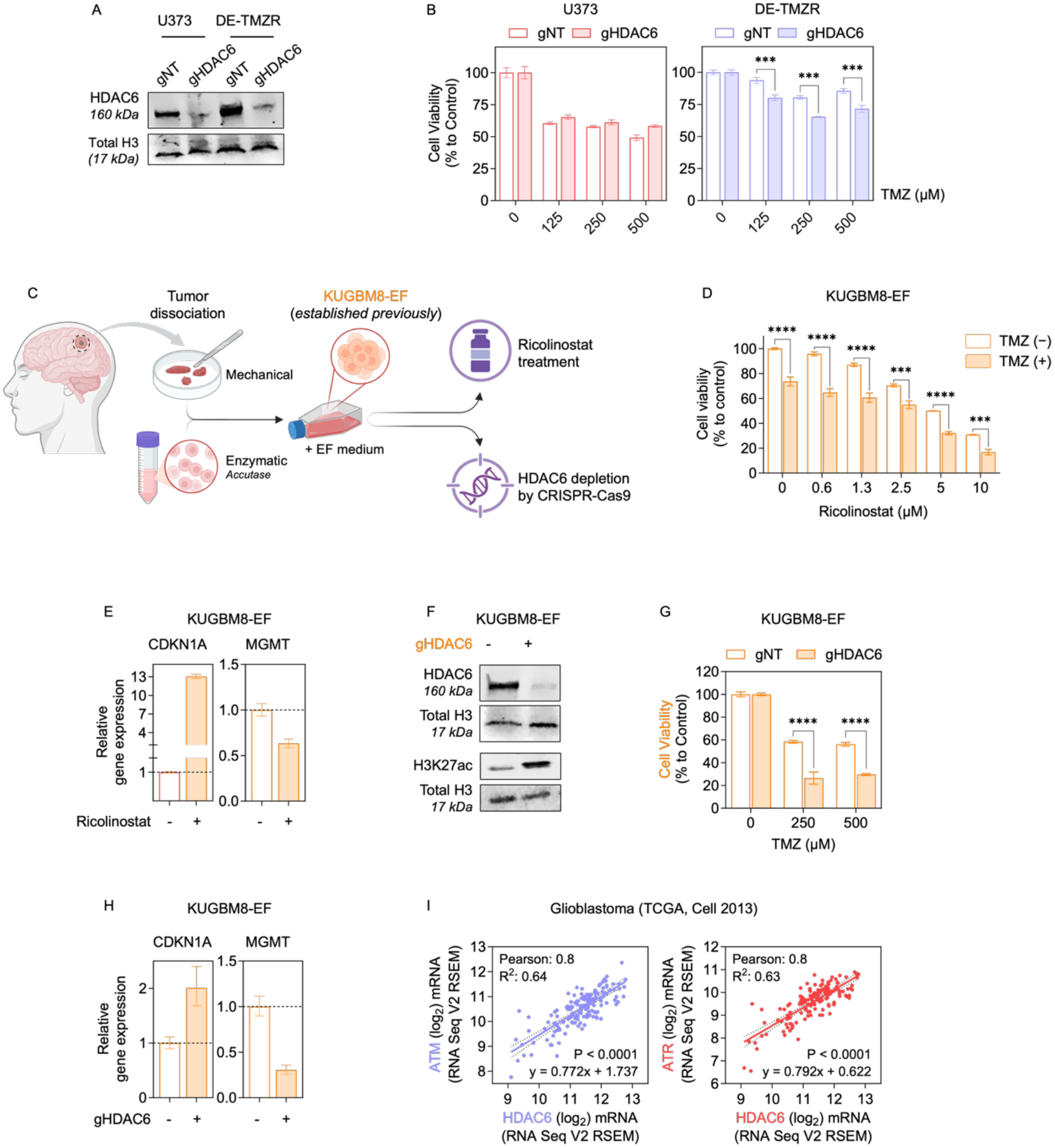
Ricolinostat treatment and HDAC6 depletion sensitize resistant and primary GBM cells (KUGBM8-EF) to TMZ. **A**. Western blot analysis confirming efficient HDAC6 knockout in U373 and DE-TMZR cells using two independent guide RNAs (HDAC6-g1 and HDAC6-g2). β-actin is used as a loading control. **B**. Cell viability of U373 and DE-TMZR cells after HDAC6 knockout (gHDAC6) versus non-targeting control (gNT) across increasing TMZ doses (125, 250, 500 µM; 72 h, CTG assay). Data are presented as mean ± SEM. Statistical analysis was performed using two-way ANOVA followed by Šídák’s multiple comparisons test. (***) p<0.001. **C**. Experimental scheme created with BioRender. Patient tumor tissue was dissociated (mechanical + Accutase) and expanded in EF medium to derive the primary GBM line KUGBM8-EF, previously established by our group ^25^. Cells were then treated with Ricolinostat or subjected to CRISPR-Cas9 mediated HDAC6 depletion. **D**. CTG results of KUGBM8-EF exposed to increasing concentrations of Ricolinostat in the absence (™) or presence (+) of TMZ (250 µM) for 72 h. Data are presented as mean ± SEM. Statistical analysis was performed using two-way ANOVA followed by Šídák’s multiple comparisons test. (***) p<0.001, (****) p<0.0001. **E**. Gene expression levels of *CDKN1A* and *MGMT* on KUGBM8-EF cells after Ricolinostat (10 µM) treatment. **F**. Western blot images representing HDAC6 and H3K27ac levels in KUGBM8-EF cells upon HDAC6 KO. **G**. CTG results of HDAC6 KO KUGBM8-EF cells in the increasing dose of TMZ for 72 h. **H**. Gene expression levels of *CDKN1A* and *MGMT* on KUGBM8-EF cells upon HDAC6 KO. **I**. Scatter plots showing correlations between HDAC6 mRNA and *ATM* (left) or *ATR* (right) mRNA in GBM (TCGA, Cell 2013).

In this study, epigenetic vulnerabilities in GBM were identified to address the ongoing challenge of TMZ resistance in clinical settings. Epigenetic drug screening highlighted HDACs, and both Ricolinostat treatment and HDAC6 knockout improved TMZ response in resistant cells as well as primary cells. Mechanistically, HDAC6 inhibition with Ricolinostat increased global acetylation levels and was associated with p21 induction, while simultaneously suppressing *MGMT* expression creating a chromatin state less permissive to DNA repair and more responsive to TMZ-induced damage. The key findings of the study are illustrated in the graphical abstract shown in **Fig. 6**.

**Figure 6.**
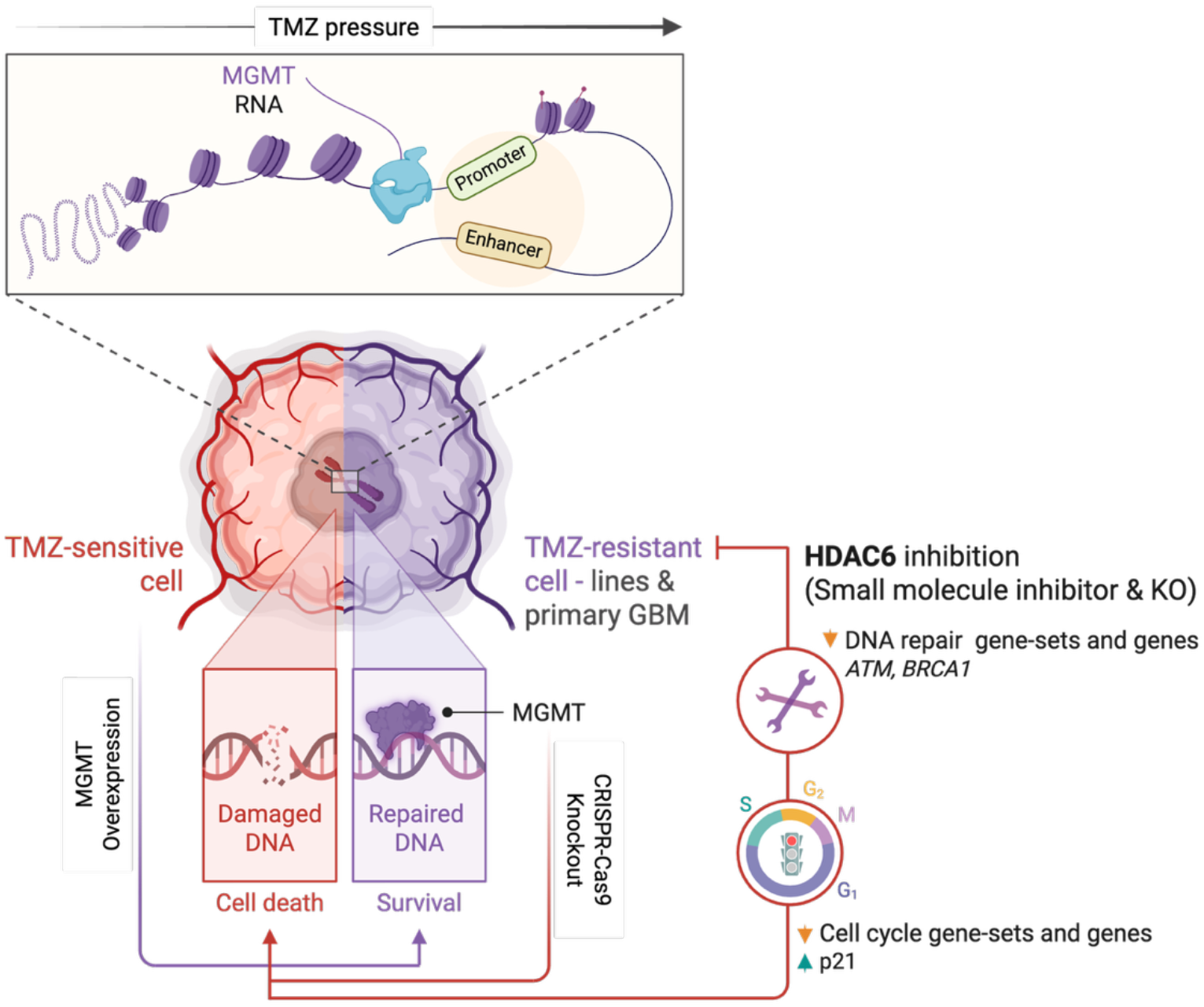
The graphical abstract of this study was created using BioRender.

## DISCUSSION

TMZ resistance remains a central barrier to effective GBM treatment, driven by both intrinsic cellular features and adaptive responses that emerge under therapeutic pressure. To interrogate these axes, we derived two TMZ-resistant models from the parental U373 line: DE-TMZR, generated by stepwise dose escalation to model adaptive/acquired reprogramming, and TMZR, established via high-dose pulse selection to enrich pre-existing, intrinsically tolerant clones. By generating two distinct resistant derivatives from the U373 parental line, we were able to dissect the divergent biological routes through which GBM evades TMZ cytotoxicity. This distinction is clinically relevant, as recurrent GBM often displays mixed resistance states shaped by evolutionary pressures and tumor heterogeneity^28^. Both resistant models showed increased TMZ tolerance relative to U373 parental, as demonstrated by a shifted dose-response with a higher IC50 and by long-term clonogenic survival. Unlike U373 parental, γH2AX was not induced in DE-TMZR or TMZR after TMZ, indicating that TMZ fails to engage its canonical damage-response program in these models. Moreover, resistant populations fail to undergo canonical G2/M arrest upon TMZ treatment, collectively mirroring features observed in recurrent GBM. These phenotypes were characterized in U373-derived acquired-resistance models; generalizability to additional GBM backgrounds remains to be tested.

Multi-omics profiling pinpointed MGMT as a dominant determinant of the resistant state, highlighting transcriptional upregulation with increased chromatin accessibility at the *MGMT* locus. As the transcriptomics profiling was performed on the resistant cells after selection, it cannot be resolved whether MGMT upregulation arose *de novo* under therapy pressure or from selection of pre-existing MGMT-high clones, which would prompt barcoding and lineage tracing analyses. Perturbation experiments established MGMT dependency: CRISPR-KO re-sensitized DE-TMZR/TMZR, while MGMT overexpression conferred TMZ tolerance and blunted γH2AX. Consistently, clinical datasets associated high MGMT with poorer outcomes and poor response to Temozolomide. Together, these data validate DE-TMZR and TMZR as complementary models anchored in MGMT-mediated repair and align with the broad literature establishing MGMT as a key determinant of TMZ-resistance and adverse clinical outcomes in GBM^29^. The DE-TMZR model also exhibited broader transcriptional and chromatin remodeling features, underscoring that adaptive resistance extends beyond MGMT reactivation and involves epigenetic reprogramming that reshapes DNA damage signalling, checkpoint dynamics and lineage plasticity^30–32^.

Epigenetic rewiring of the tumor cell transcriptome is an important adaptive mechanism for therapy pressure^33,34^. This plasticity enables tumor cells to reorganize chromatin accessibility, alter enhancer-promoter interactions, and modulate transcription programs that buffer against cytotoxic stress. Therefore, it is reasonable to suggest that epigenetic regulation contributes to acquired TMZ resistance. Indeed, targeted approaches have been shown to potentially overcome this resistance. For instance, Li *et al*. showed that upon EGFR activation, EZH2-mediated methylation of tonicity-responsive enhancer binding protein (NFAT5) leads to upregulation of MGMT, resulting in a poor response to TMZ. Inhibiting this methylation improved TMZ efficacy in orthotopic and patient-derived xenograft (PDX) models^35^. Moon *et al*. demonstrated that destabilization of Methyl CpG binding domain protein 3 (MBD3), a key element of the repressive nucleosome remodeling and deacetylase (NuRD) complex, can overcome TMZ-chemoresistance by promoting the neural differentiation of the GBM stem-like cell subpopulation^36^. Consistent with these findings, our RNA-seq and ATAC-seq analyses revealed pronounced transcriptional and accessibility gains at the MGMT locus and highlighted widespread alterations in DNA-repair and cell-cycle programs in resistant derivatives. These observations align with the idea that acquired resistance emerges through adaptive chromatin states, in contrast to intrinsic resistance which relies more heavily on constitutive MGMT expression.

We next asked whether the adaptive changes underlying acquired TMZ resistance introduce new epigenetic vulnerabilities that are not present in MGMT-driven intrinsic resistance. To this end, we carried out an epigenetic inhibitor screen in parental U373 cells, their resistant derivatives and T98G cells. Strikingly, while T98G cells showed no notable response to epigenetic perturbation, consistent with innate MGMT-driven resistance lacking broader epigenetic remodeling, the acquired-resistant line DE-TMZR displayed a clear epigenetic vulnerability. The selective response of DE-TMZR cells, but not T98G cells, indicates that epigenetic dependencies arise specifically through therapy-induced adaptations rather than through MGMT-driven intrinsic resistance. Notably, PARP inhibitors also potentiated TMZ activity, consistent with known synthetic lethal interactions between alkylating damage and PARP-mediated base-excision repair^37^, further reinforcing the connection between TMZ pressure and DNA-repair compensation.

Among epigenetic classes, HDAC inhibitors were clustered, and the HDAC6 inhibitor Ricolinostat increased apoptosis, evidenced by cleaved PARP and elevated histone H3 acetylation, consistent with on-target HDAC6 inhibition. Previous reports implicate HDAC6 in GBM biology, including proliferation, invasion, stress responses, and DNA-damage programs^38–41^, and selective HDAC6 inhibitors have been noted to potentiate TMZ in preclinical models; our findings are consistent with, and extend, these observations by demonstrating activity in parental as well as in resistant derivatives, with effects more pronounced in the resistant state.

Beyond the histone-acetylation readouts presented here, HDAC6 is predominantly cytosolic and its canonical substrates lie outside the histone code. HDAC6 deacetylates α-tubulin to regulate microtubule dynamics and intracellular trafficking^42^, deacetylates HSP90 to license chaperoning of multiple client proteins, and, through its zinc-finger ubiquitin-binding domain, orchestrates aggresome formation and the autophagic clearance of misfolded protein aggregates^43,44^. The increase in histone H3 acetylation we observe upon Ricolinostat treatment is therefore unlikely to reflect direct deacetylation of histones by HDAC6 itself; rather, it is consistent with engagement of class I HDAC activity, given that Ricolinostat (ACY-1215) is preferentially HDAC6-selective but loses isoform selectivity at the micromolar concentrations used in cellular assays^45^. Importantly, the cytosolic functions of HDAC6 intersect directly with proteostasis of DNA-damage-response factors via HSP90 chaperoning^44^, offering a non-histone mechanism that complements the transcriptional changes we report and could contribute to the loss of DNA-repair capacity and the restoration of TMZ sensitivity observed upon HDAC6 inhibition or knockout. Future studies dissecting the relative contributions of nuclear (histone, transcriptional) and cytosolic (microtubule, aggresome and HSP90-client) effects of HDAC6 inhibition in TMZ-resistant GBM will help refine the therapeutic mechanism.

Several HDAC inhibitors, such as Vorinostat, Panobinostat, Romidepsin and Belinostat, are FDA-approved for hematologic malignancies, yet none is approved for GBM, underscoring the need to define where HDAC targeting adds value in this setting^46^. Motivated by the screen results and on-target engagement with the HDAC6-targeting agent Ricolinostat, we next characterized transcriptome-level changes after Ricolinostat to map pathways linked to chemosensitivity. Across U373, DE-TMZR, and TMZR, Ricolinostat suppressed cell-cycle and DNA-repair programs, highlighted by shared downregulated genes, enrichment networks, and a chord diagram converging on repair terms, with ATM protein also reduced, and Hallmark analyses showing concordant decreases in mitotic spindle, DNA repair, G2/M checkpoint, E2F targets, and MYC targets. Critically, several key DNA-repair regulators, including *ATM, CHK1, PMS2, BRCA1* and *RAD51* were significantly downregulated at the transcript levels, with ATM suppression also confirmed at the protein level. Functionally, these transcriptional changes translated into a pronounced G2/M arrest accompanied by a robust upregulation of p21 (*CDKN1A*) across all models, enforcing checkpoint activation and preventing progression through the cell cycle in the presence of DNA lesions. Together, the combined loss of ATM-mediated repair capacity, downregulation of additional HR and MMR components, and induction of p21-driven cell-cycle arrest provides a mechanistic basis for the enhanced TMZ sensitivity observed upon HDAC6 inhibition, positioning HDAC6 as a critical regulator of the dual repair and proliferative programs that TMZ-resistant GBM cells depend on for survival.

To validate that HDAC6 represents a genuine vulnerability in TMZ-resistant GBM, we next evaluated both genetic and pharmacologic perturbation of HDAC6 in established cell lines and patient-derived models. Initial siRNA screening showed HDAC6 depletion reproducibly sensitized resistant cells to TMZ, prompting the generation of CRISPR-mediated HDAC6 KO lines. In the primary GBM line KUGBM8-EF, Ricolinostat enhanced TMZ responses and modulated *CDKN1A* and *MGMT* levels, while HDAC6 loss increased H3K27ac, reduced *MGMT*, and restored TMZ sensitivity. TCGA correlations between HDAC6 and *ATM/ATR* (Fig. 5I), together with transcriptomic downregulation of key elements of DDR, further support HDAC6 as a regulator of DNA-damage signaling. Overall, these findings demonstrate that HDAC6 inhibition disrupts chromatin state and DNA-repair capacity and re-sensitizes diverse GBM models to TMZ.

From a clinical perspective, our results provide a strong rationale for exploring HDAC6 inhibitors as combinatorial partners with TMZ, particularly in recurrent GBM where MGMT-driven resistance, therapy-induced epigenetic plasticity, and DDR pathway upregulation are prevalent. While several pan-HDAC inhibitors have been evaluated clinically, their CNS penetration and toxicity profiles have limited translation to GBM^47^. HDAC6-selective inhibitors, which exhibit improved tolerability, may offer a more feasible therapeutic approach, especially in a setting requiring prolonged treatment or combination with cytotoxic agents. Our findings further suggest that HDAC6 inhibition could potentially reduce the threshold of TMZ required for cytotoxicity, which may help mitigate systemic toxicity.

In conclusion, our work identifies HDAC6 as a key epigenetic dependency that emerges specifically during the acquisition of TMZ resistance. By integrating chromatin accessibility, transcriptional reprogramming, cell-cycle control, and DNA-repair capacity, HDAC6 enables GBM cells to survive TMZ-induced damage. Targeting HDAC6 disrupts these adaptive programs, restoring sensitivity to TMZ across both established and primary models. These findings provide a compelling mechanistic rationale for the development of HDAC6-directed combination regimens and support further translational efforts to evaluate HDAC6 inhibitors in recurrent, treatment-experienced GBM.

## AUTHOR CONTRIBUTIONS

Study design: F.S.A., T.B.O.; Methodology: F.S.A., A.P.C., U.O., T.B.O.; Data generation and analysis: F.S.A., B.C., N.D., E.Y.K., D.U., M.P., A.P.C., J.D.; Data interpretation: F.S.A., B.C., N.D., U.O., C.A., T.B.O.; Drafted the manuscript: F.S.A., B.C., N.D., T.B.O.; Approved final manuscript: all authors.

## ACKNOWLEDGEMENTS

The authors gratefully acknowledge the use of the services and facilities of the Koç University Research Center for Translational Medicine (KUTTAM). We also thank Dr. Ilknur Sur-Erdem, Dr. Ezgi Özyerli-Göknar, Dr. Alişan Kayabölen, and Dr. Tamer Önder for their valuable support and scientific input.

## FUNDING

This work was supported in part by the EMBO Short-Term Fellowship (to F.S.A.), TÜBİTAK 1003 grant #216S461 (to T.B.O.), and EU COST Action NET4BRAIN (grant CA22103). The funders had no role in study design, data collection and analysis, decision to publish, or preparation of the manuscript.

## COMPETING INTERESTS

M.P., A.P.C. and U.O. are co-founders of Caeruleus Genomics Ltd (Entelo Bio) and inventors on several patents related to sequencing technologies filed by Oxford University Innovations. The other authors declare no competing interests.

## DATA AND CODE AVAILABILITY

RNA-sequencing and ATAC-sequencing data have been deposited in the EMBL-EBI ArrayExpress database under accession numbers E-MTAB-16087 and E-MTAB-16098. Figures created with BioRender.com have been licensed for publication (Fig. 1A: EZ290UUFS6; Fig. 1H: GO290UV1RJ; Fig. 2H: YW290V0K1V; Fig. 3B: YO290V0T8G; Fig. 5C: ZT290V1A9Q; Fig. 6: AG290V1HTT; Fig. S1A: GX290V1RFT; Fig. S2A: EG290V24Q1).

## Notes

### Competing Interest Statement

MP, A.P.C. and U.O are co-founders of Caeruleus Genomics Ltd (Entelo Bio) and inventors on several patents related to sequencing technologies filed by Oxford University Innovations.

