## Supplementary Files for "Chemical screens identify HDAC6 as an epigenetic vulnerability in acquired Temozolomide-resistant models of glioblastoma"

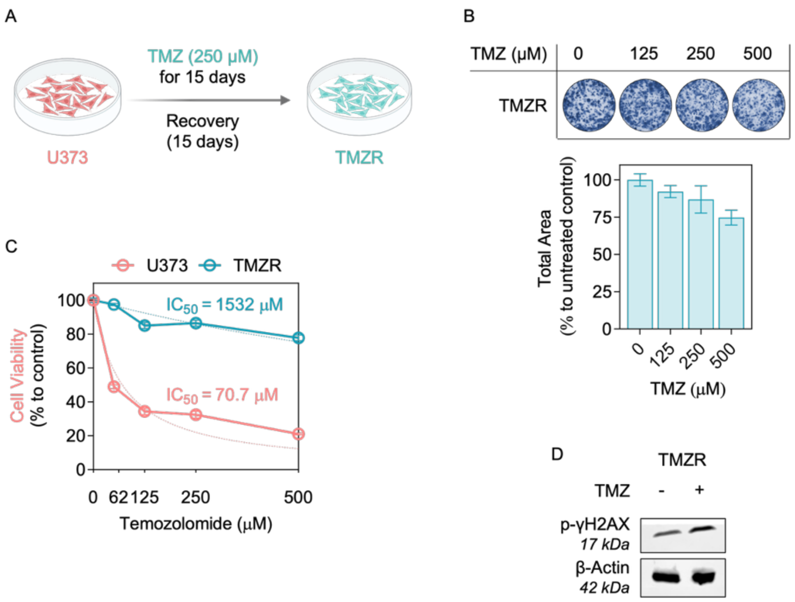


**Fig. S1. TMZ-resistance induction scheme for TMZR.**

- **A.** Schematics of the establishment of TMZ-resistant U373 cells (TMZR) by fixed-dose selection. Cells were exposed to 250 µM TMZ for 15 days, followed by a 15-day drug-free recovery period. Created with BioRender.
- **B.** Cells were treated with TMZ for 48 h, and after drug removal, they were allowed to proliferate for 10 days. Colony areas were measured and normalized to the respective controls.
- **C.** CTG cell viability results were obtained after 5 days of TMZ treatment.
- **D.** γ-H2AX protein levels were assessed by Western blot analysis after TMZ treatment (250 µM, 24 h); β-actin is used as a loading control.

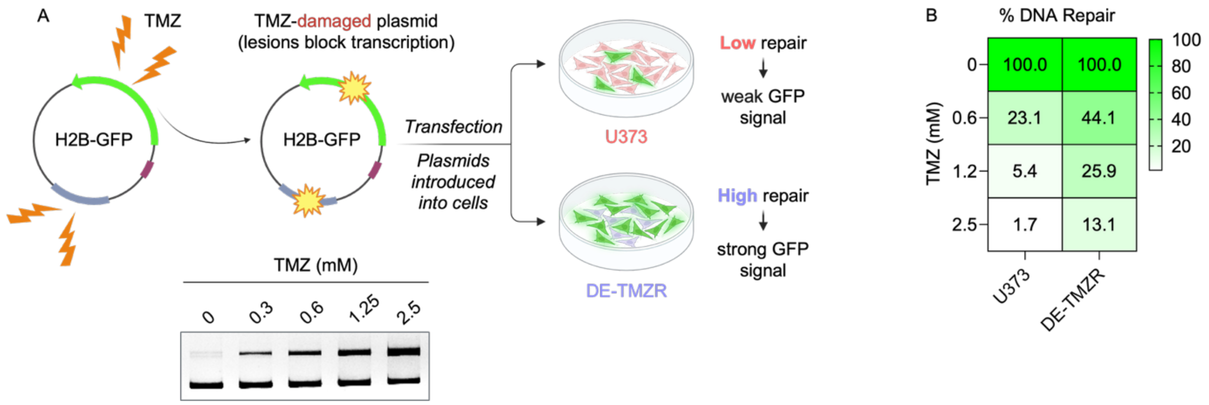


**Fig. S2. Host cell reactivation (HCR) assay for evaluating DNA-repair capacities of U373 and DE-TMZR cells.**

**A.** The schematic for the HCR assay. Plasmids are damaged by TMZ treatment. Both damaged and undamaged plasmids are then transfected into parental U373 and DE-TMZR cells.

**B.** Quantification of intracellular GFP signal 72 h after transfection, measured by flow cytometry and shown as a heat map (% GFP-positive cells relative to undamaged control).

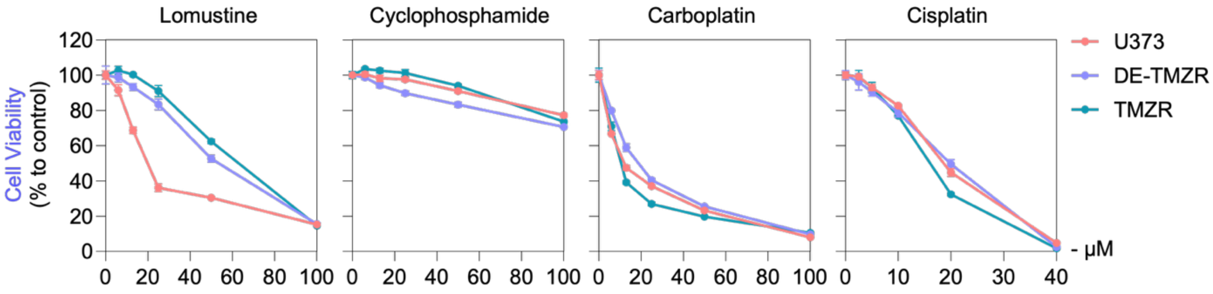


**Fig. S3. Cell viability was evaluated for the alkylating chemotherapeutic agents Lomustine, Cyclophosphamide, Carboplatin, and Cisplatin.** The cells were exposed to the indicated doses for 72 h, and viability was determined using the CTG assay. Statistical significance was assessed by two-way ANOVA followed by Šídák's multiple comparisons test. *Asterisks indicate significance ($p < 0.0001$).

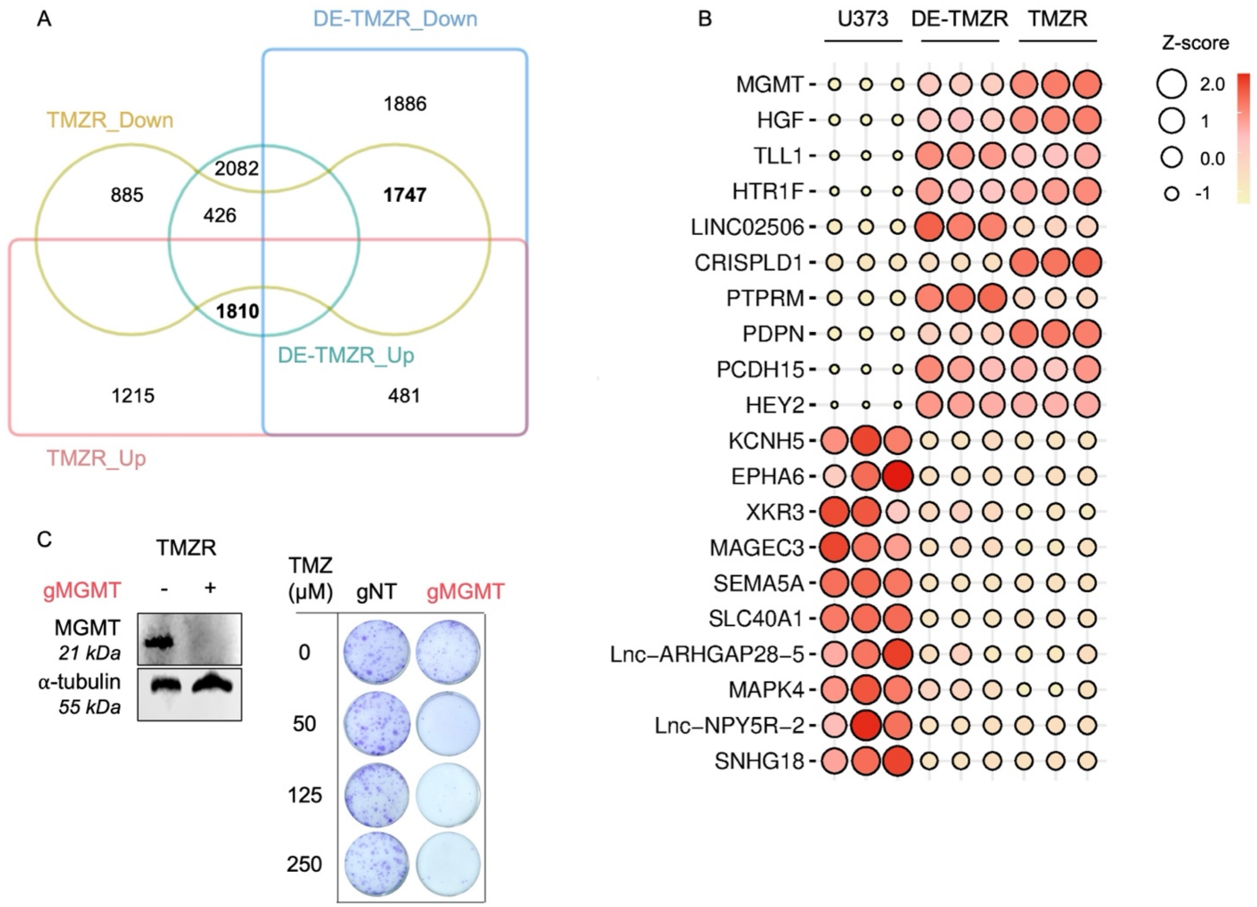


**Fig. S4. Transcriptomic profiling of TMZR cells and functional validation of MGMT.**

- **A.** Venn diagram showing overlaps among RNA-seq DEGs ($p \leq 0.05$) from TMZR and DE-TMZR cells compared with parental U373, separated into up- and downregulated sets.
- **B.** Balloon plot of genes from the shared DEGs in (A), common upregulated (n=1810), and downregulated (n=1747) genes in TMZR and DE-TMZR compared with parental U373.
- **C.** MGMT protein levels and clonogenic assay results of TMZR cells following MGMT knockout, with representative colony images and quantifications.

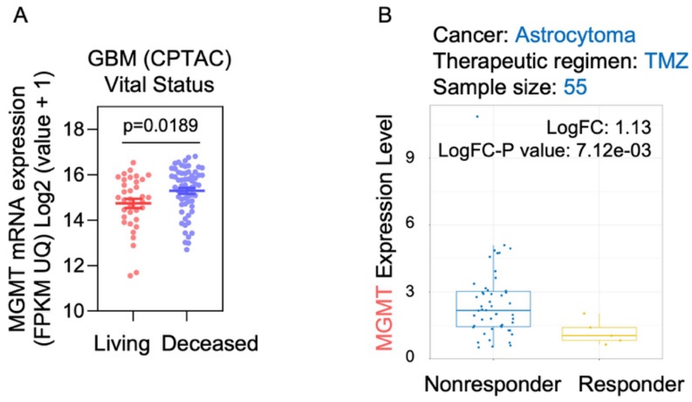


**Fig. S5. Clinical datasets supporting the association between *MGMT* expression and poor TMZ response.**

- **A.** CPTAC dataset analysis showing that *MGMT* mRNA levels are significantly higher in deceased GBM patients compared with living cases (p = 0.0189).
- **B.** CTR-DB dataset analysis of astrocytoma patients (n = 55) receiving TMZ treatment, indicating that high *MGMT* expression is associated with non-responder status (LogFC = 1.13, p = $7.12 \times 10^{-3}$).

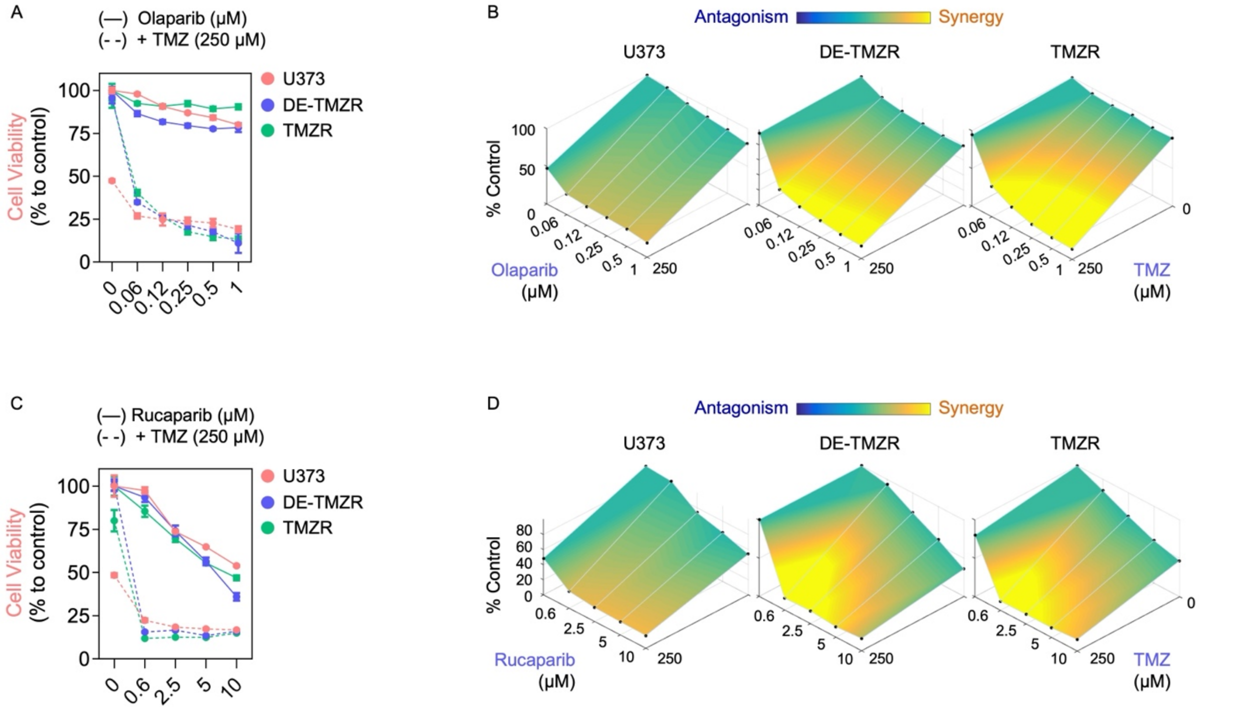


**Fig. S6. Proof-of-principle of TMZ-PARP inhibitor synergy in parental and TMZ-resistant GBM cells.**

- **A.** Cell viability of parental (U373) and TMZ-resistant (DE-TMZR, TMZR) cells treated with increasing concentrations of Olaparib (solid lines) or Olaparib + TMZ (250 µM) (dashed lines) for 72 h, expressed as % of untreated control.
- **B.** 3D surface synergy plots generated using Combenefit**,** showing the interaction between Olaparib and TMZ in U373, DE-TMZR, and TMZR cells. Color scale indicates drug interaction, from antagonism (blue) to synergy (yellow).
- **C.** Cell viability curves for Rucaparib (solid lines) and Rucaparib + TMZ (250 µM) (dashed lines) under the same conditions as in (A).
- **D.** Corresponding 3D synergy surface plots illustrating the combined effects of Rucaparib and TMZ.

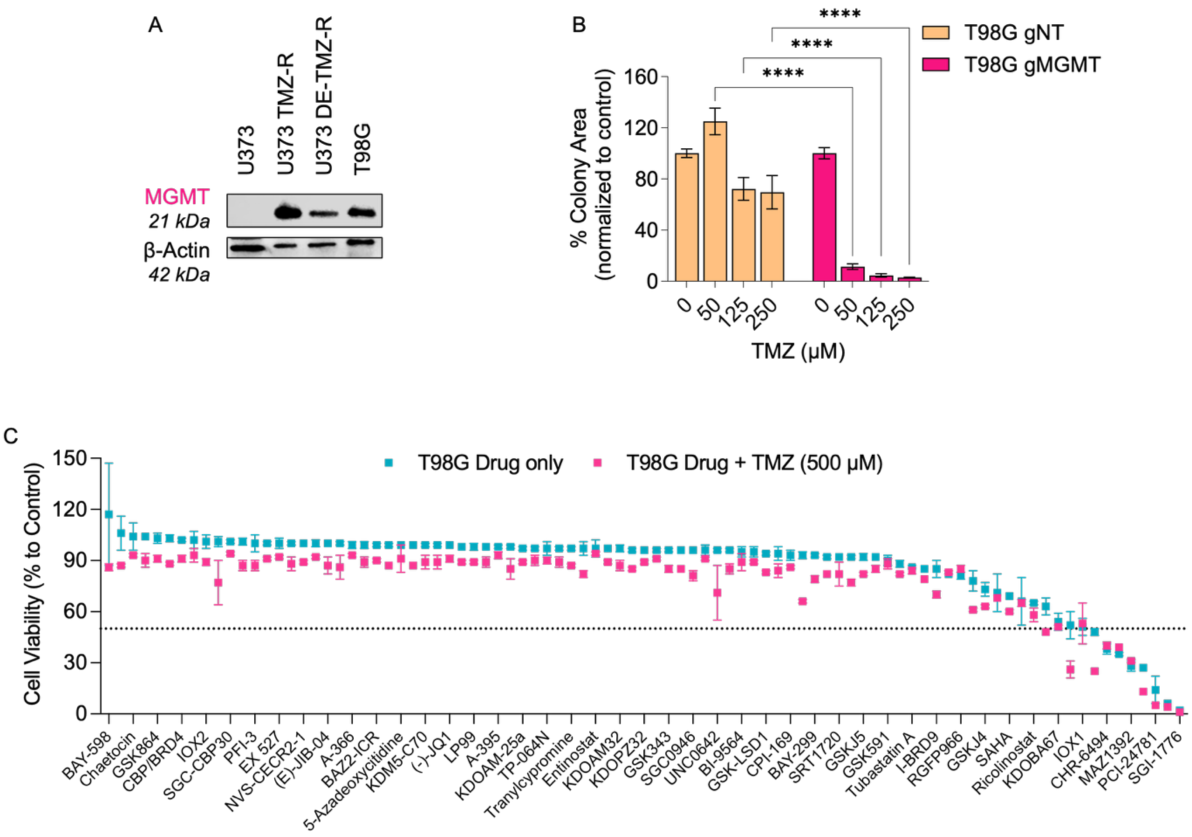


**Fig. S7. MGMT status and an epigenetic inhibitor screen in T98G cells.**

- **A.** MGMT protein levels were assessed by Western blot analysis in U373, U373-TMZ-R, U373-DE-TMZ-R, and T98G cells; β-actin was used as a loading control.
- **B.** Clonogenic survival of T98G cells following MGMT knockout (gMGMT); gNT denotes the non-targeting guide control.
- **C.** Cell viability results from screens performed on T98G cells. Cells were treated with the respective drug, with or without TMZ (500 µM) for 72 h. Cell viability was measured using the CTG assay and normalized to untreated controls.

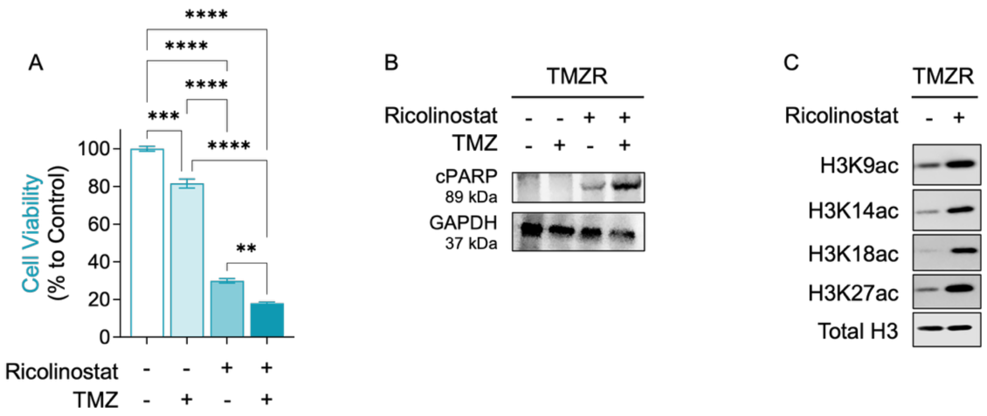


**Fig. S8. Ricolinostat induces apoptosis and increases histone acetylation in TMZR cells.**

- **A.** Cell viability results of the combination of Ricolinostat (10 µM) and TMZ (250 µM). Statistical analysis was performed using one-way ANOVA followed by Tukey's multiple comparisons test. (**) p<0.01, (***) p<0.001, (****) p<0.0001.
- **B.** Western blot showing PARP cleavage following 36 h treatment with Ricolinostat (10 µM) and/or TMZ (250 µM), with GAPDH used as a loading control.
- **C.** Western blot analysis of Histone 3 acetylation marks after 36 h of Ricolinostat (10 µM) treatment, with total H3 as a loading control.

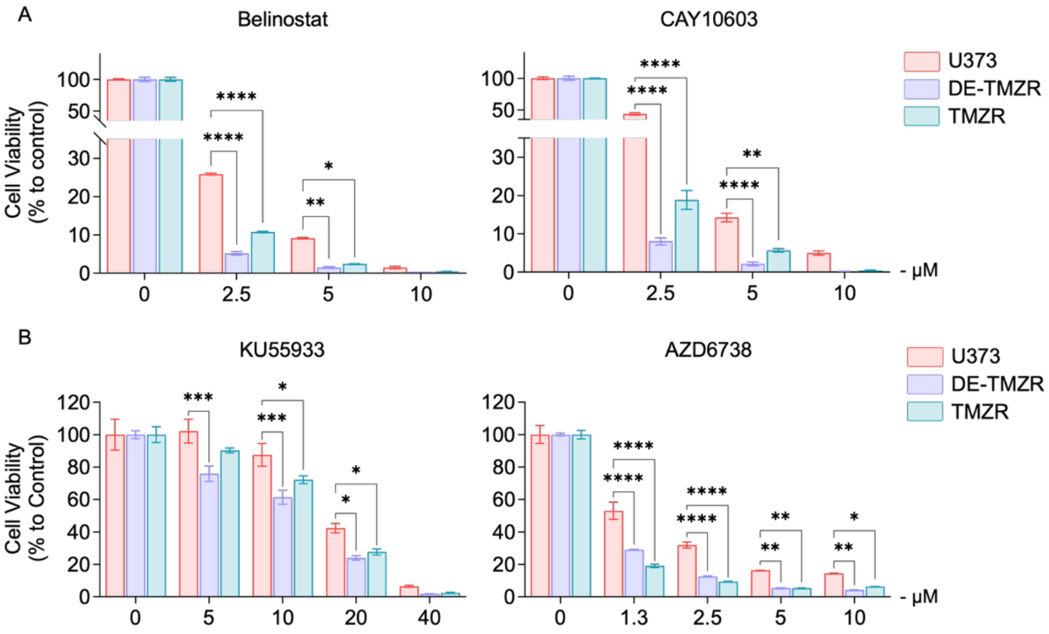


**Fig. S9. TMZ-resistant cells exhibit increased vulnerability to HDAC6 and ATM/ATR inhibitors.**

- **A.** Cell viability results for the HDAC6 inhibitors Belinostat and CAY10603 in parental U373 and TMZ resistant (DE-TMZR, TMZR) cells were obtained using the CTG assay after 72 h. Statistical analysis was performed using two-way ANOVA followed by Šídák's multiple comparisons test. (**) $p<0.01$, (***) $p<0.001$, (****) $p<0.0001$.
- **B.** Cell viability results for the DDR inhibitors KU55933 (ATM) and AZD6738 (ATR) in parental U373 and TMZ resistant (DE-TMZR, TMZR) cells were obtained using the CTG assay after 72 h. Statistical analysis was performed using two-way ANOVA followed by Dunnett's multiple comparisons test. (*) $p<0.05$, (**) $p<0.01$, (***) $p<0.001$, (****) $p<0.0001$.

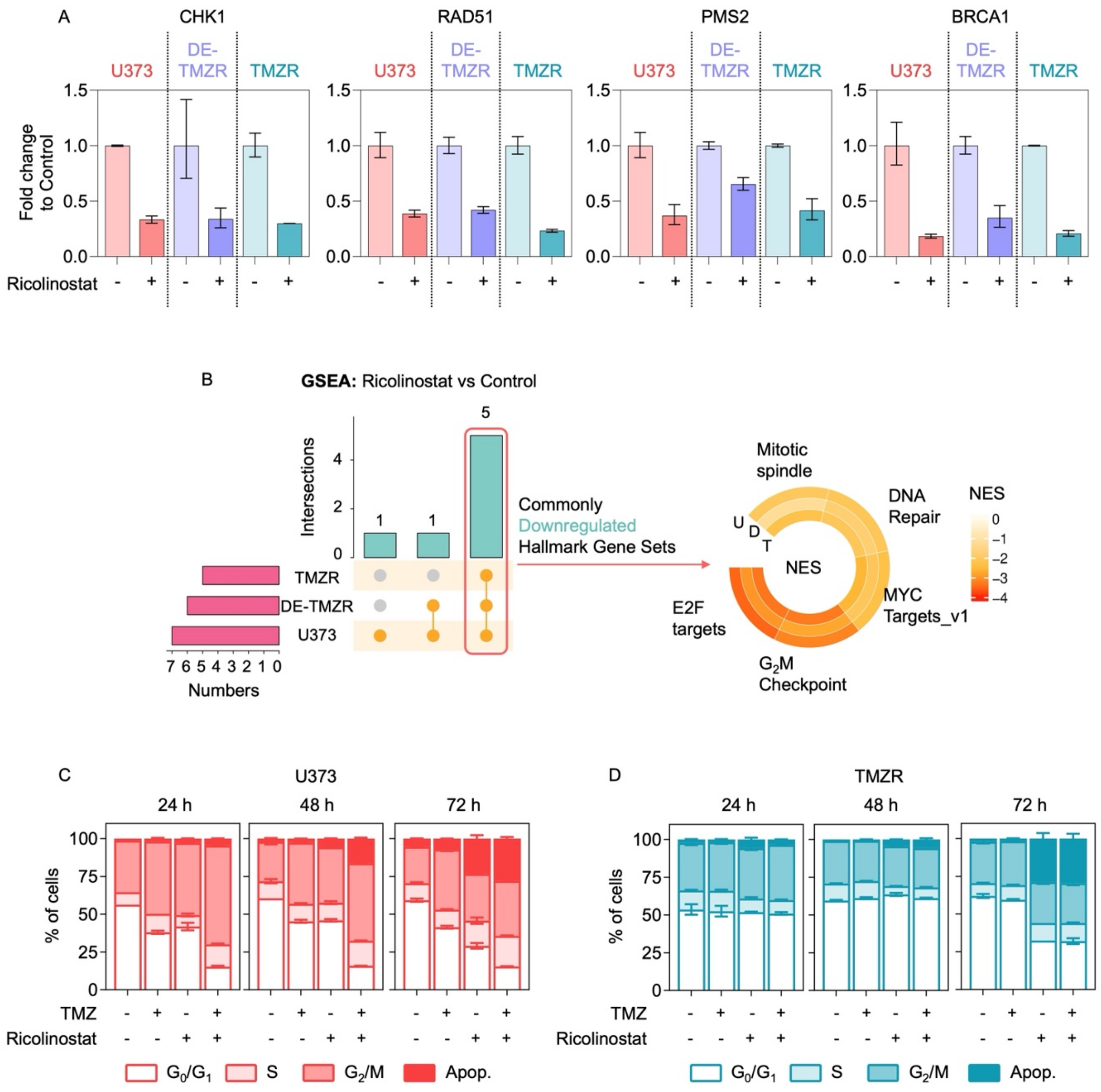


**Fig. S10. Ricolinostat downregulates Hallmark cell-cycle and DNA repair programs.**

**A.** RT-qPCR validation of genes found to change 24 h after Ricolinostat across U373, DE-TMZR, and TMZR.

**B.** Gene Set Enrichment Analysis (GSEA; MSigDB Hallmark collection) of Ricolinostat-treated cells across parental U373 and two TMZ-resistant derivatives (DE-TMZR and TMZR). Common overlaps among Hallmark gene sets (five in total) are shown by the UpSet plot (left). The corresponding normalized enrichment scores (NES) for these commonly downregulated pathways, including mitotic spindle, DNA repair, MYC targets v1, G2/M checkpoint, and E2F targets, are visualized in the heatmap on the right. Analyses were performed using a fold-change (FC) cutoff of 0.5. **U**: U373, **D**: DE-TMZR, **T**: TMZR.

**C.** Time-course cell-cycle profiling after Ricolinostat (10 µM) ± TMZ (250 µM) in U373 cells.

**D.** Time-course cell-cycle profiling after Ricolinostat (10 µM) ± TMZ (250 µM) in TMZR cells.

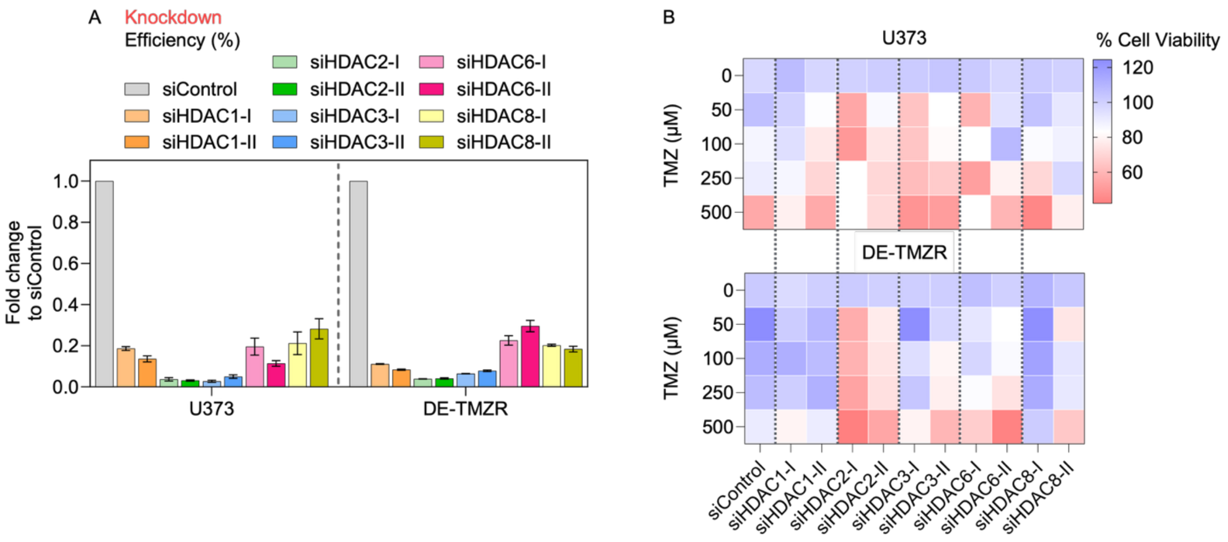


**Fig. S11. HDAC depletion was achieved through siRNA-mediated targeting, followed by assessment of TMZ response.**

**A.** RT-qPCR confirmed gene-level knockdown of *HDAC1, HDAC2, HDAC3, HDAC6,* and *HDAC8*, normalized to siControl.

**B.** Heatmaps showing cell viability (%) measured by CTG assay (72 h) across increasing TMZ concentrations following siRNA knockdown in U373 (top) and DE-TMZR (bottom); values are normalized to each siRNA's untreated control.

**Table S1.** List of 96 epigenetic probes with working concentrations (µM)

and targets in the drug library utilized in the study.

| Class | Compound | Target | Concentration |
|---|---|---|---|
| **Arginine methyltransferase** | SGC707 | PRMT3 | 1 |
| | MS049 | PRMT4/6 | 1 |
| | MS023 | PRMT4/6 | 1 |
| | GSK591 | PRMT5 | 1 |
| | MS409N | PRMT4/6 - inactive control | 1 |
| | TP-064 | PRMT4 | 1 |
| | TP-064N | PRMT4 - inactive control | 1 |
| | AMI-1 | PRMT | 50 |
| **Bromodomain** | (+)-JQ1 | BRD2,BRD3,BRD4,BRDT (BET) | 1 |
| | (-)-JQ1 | Negative Control | 1 |
| | PFI-1 | BRD2,BRD3,BRD4,BRDT (BET) | 5 |
| | I-BET | BRD2,BRD3,BRD4 | 1 |
| | Bromosporine | pan-Bromodomain | 1 |
| | CBP/BRD4 | CBP,BRD4 | 1 |
| | SGC-CBP30 | CREBBP, EP300 | 1 |
| | I-CBP112 | CREBBP, EP300 | 1 |
| | RVX-208 | BRD2,BRD3,BRD4,BRDT (BET) | 5 |
| | SMARCA | SMARCA, PB1 | 2,5 |
| | PFI-3 | SMARCA2/4, PB1(5) | 1 |
| | GSK2801 | BAZ2A,BAZ2B | 1 |
| | PFI-4 | BRPF1B | 1 |
| | OF-1 | pan-BRPF | 5 |
| | BAZ2-ICR | BAZ2A,BAZ2B | 1 |
| | NI-57 | pan-BRPF | 1 |
| | LP99 | BRD97/9 | 1 |
| | BI-9564 | BRD7/9 | 1 |
| | NVS-CECR2-1 | CECR2 | 1 |
| | GSK8814 | ATAD2 | 10 |

| | | | |
|---|---|---|---|
| | GSK8815 | ATAD2 - inactive control | 10 |
| | BAY-299 | BRD1, TAF1 | 1 |
| | I-BRD9 | BRD9 | 10 |
| | TP-472 | BRD9 | 1 |
| | TP-472N | BRD9 - inactive control | 1 |
| **Dehydrogenase** | GSK864 | IDH | 5 |
| **DNA Methyltransferase** | 5-Azacitidine | DNMT | 10 |
| | 5-Azadeoxycitidine | DNMT1/3 | 5 |
| **Halofuginol** | MAZ1805 | inactive control | 1 |
| | MAZ1392 | | 1 |
| **Histone acetyltransferase** | C646 | p300/CBP | 1 |
| **Histone deacetylase** | Belinostat | Hydroxamic acids | 5 |
| | CXD101 | HDAC | 1 |
| | Valproic acid | Aliphatic acid compounds | 1000 |
| | Entinostat | Ortho-amino anilides | 0,5 |
| | SAHA | Hydroxamic acids | 2,5 |
| | TrichostatinA | Hydroxamic acids | 0,5 |
| | SRT1720 | SIRT1 | 1 |
| | EX527 | SIRT1 | 1 |
| | CI-994 | HDAC1/2/3 | 1 |
| | RGFP966 | HDAC3 | 10 |
| | PCI-34051 | HDAc | 5 |
| | Ricolinostat | HDAC6 | 10 |
| | Tubastatin A HCL | HDAC6 | 10 |
| | PCI-24781 | HDAC | 5 |
| | Romidepsin | HDAC | 1 |
| | Mocetinostat | HDAC | 10 |
| | Santacruzamate | HDAC2 | 50 |
| | TMP269 | HDAC4/5/7/9 | 10 |
| | AGK2 | SIRT2 | 10 |
| **Histone demethylase** | Methylstat | | 2,5 |
| | (E)-JIB-04 | pan-JmjC | 0,05 |

| | | | |
|---|---|---|---|
| **Histone methyltransferase** | CPI-360 | EZH1/2 | 10 |
| | UNC0638 | G9a,GLP | 1 |
| | UNC0642 | G9a,GLP | 1 |
| | A-366 | G9a,GLP | 2 |
| | Chaetocin | SUV39H1 | 0,05 |
| | PFI-2 | SETD7 | 2 |
| | SGC0946 | DOT1L | 7,5 |
| | GSK343 | EZH2 | 3 |
| | UNC1999 | EZH2 | 1 |
| | LLY-507 | SMYD2 | 1 |
| | A-196 | SUV420H1/H2 | 1 |
| | BAY-598 | SMYD2 | 1 |
| | CPI-169 | EZH1/2 | 10 |
| | UNC2400 | EZH2 - inactive control | 1 |
| **Kinase Inhibitor** | K00135 | ATP competitive- PIM | 1 |
| | 5-Iodotubercidin | ATP mimetic-Haspin | 1 |
| | SGI-1776 | Haspin | 10 |
| | CHR-6494 | Haspin | 1 |
| **Lysine demethylase** | Tranylcypromine | LSD1 | 20 |
| | GSK-LSD1 | LSD1 | 0,5 |
| | GSKJ4 | JMJD3, UTX, JARID1B | 10 |
| | GSKJ5 | inactive control | 10 |
| | IOX1 | pan-2-OG | 40 |
| | KDOBA67 | KDM6A/B | 10 |
| | KDOAM-25a | JARID | 1 |
| | KDM5-C70 | JARID1 | 10 |
| | KDOAM32 | JARID | 1 |
| | KDOPZ-32a | KDM5 | 1 |
| | KDOOA012000 | KDM2 | 1 |
| **Methyl Lysine Binder** | OICR-9429 | WDR5 | 1 |
| | UNC1215 | L3MBTL3 | 5 |
| | A-395 | EED | 1 |

| | | | |
|---|---|---|---|
| | A-395N | EED- inactive control | 1 |
| **Peptidyl arginine deiminase** | GSK484 | PAD4 | 1 |
| | GSK106 | PAD4 | 1 |
| **Poly ADP ribose polymerase** | Olaparib | PARP | 10 |
| | Rucaparib | PARP | 10 |
| **Prolyl-Hydroxylase** | IOX2 | PHD2 | 10 |

**Table S2.** Adapters used in ATAC-seq

| Adapter ID | Sequence |
|---|---|
| Ad1 | AATGATACGGCGACCACCGAGATCTACACTCGTCGGCAGCGTCAGATGTG |
| Ad2 | CAAGCAGAAGACGGCATACGAGATTCGCCTTAGTCTCGTGGGCTCGGAGATGT |
| Ad3 | CAAGCAGAAGACGGCATACGAGATCTAGTACGGTCTCGTGGGCTCGGAGATGT |
| Ad4 | CAAGCAGAAGACGGCATACGAGATTTCTGCCTGTCTCGTGGGCTCGGAGATGT |
| Ad5 | CAAGCAGAAGACGGCATACGAGATGCTCAGGAGTCTCGTGGGCTCGGAGATGT |
| Ad6 | CAAGCAGAAGACGGCATACGAGATAGGAGTCCGTCTCGTGGGCTCGGAGATGT |
| Ad7 | CAAGCAGAAGACGGCATACGAGATCATGCCTAGTCTCGTGGGCTCGGAGATGT |
| Ad8 | CAAGCAGAAGACGGCATACGAGATGTAGAGAGGTCTCGTGGGCTCGGAGATGT |
| Ad9 | CAAGCAGAAGACGGCATACGAGATCAGCCTCGGTCTCGTGGGCTCGGAGATGT |
| Ad10 | CAAGCAGAAGACGGCATACGAGATTGCCTCTTGTCTCGTGGGCTCGGAGATGT |
| Ad11 | CAAGCAGAAGACGGCATACGAGATTCCTCTACGTCTCGTGGGCTCGGAGATGT |
| Ad12 | CAAGCAGAAGACGGCATACGAGATTCATGAGCGTCTCGTGGGCTCGGAGATGT |
| Ad13 | CAAGCAGAAGACGGCATACGAGATCCTGAGATGTCTCGTGGGCTCGGAGATGT |
| Ad14 | CAAGCAGAAGACGGCATACGAGATTAGCGAGTGTCTCGTGGGCTCGGAGATGT |
| Ad15 | CAAGCAGAAGACGGCATACGAGATGTAGCTCCGTCTCGTGGGCTCGGAGATGT |
| Ad16 | CAAGCAGAAGACGGCATACGAGATTACTACGCGTCTCGTGGGCTCGGAGATGT |
| Ad17 | CAAGCAGAAGACGGCATACGAGATAGGCTCCGGTCTCGTGGGCTCGGAGATGT |
| Ad18 | CAAGCAGAAGACGGCATACGAGATGCAGCGTAGTCTCGTGGGCTCGGAGATGT |
| Ad19 | CAAGCAGAAGACGGCATACGAGATCTGCGCATGTCTCGTGGGCTCGGAGATGT |

**Table S3.** Sequences of oligonucleotides used in this study.

| Gene | Forward primer sequence | Reverse primer sequence |
|---|---|---|
| **RT-qPCR** | | |
| ATM | CATCGCATGTGATTAAAGCA | TTCTGATAGGAATCAGGGC |
| ATR | CAATTGTGGAGGAGATTTCC | CTTCTGAGAACTCTTGTATCTG |
| BRCA1 | CTGAAGACTGCTCAGGGCTATC | AGGGTAGCTGTTAGAAGGCTGG |
| CHK1 | CGGTATAATAATCGTGAGCG | TTCCAAGGGTTGAGGTATGT |
| GAPDH | AGCCACATCGCTCAGACAC | GCCCAATACGACCAAATCC |
| HDAC1 | CATCTCCTCAGCATTGGCTT | CGAATCCGCATGACTCATAA |
| HDAC10 | GAACAGCCACATCCAGGG | CCTCTTAGATGGGATGCTGG |
| HDAC2 | ATGAGGCTTCATGGGATGAC | ATGGCGTACAGTCAAGGAGG |
| HDAC3 | CTGTGTAACGCGAGCAGAAC | GCAAGGCTTCACCAAGAGTC |
| HDAC4 | CTGGTCTCGGCCAGAAAGT | CGTGGAAATTTTGAGCCATT |
| HDAC5 | GAACTGGGCATGGCTCTTG | GGGAACCATCCTTGGAAATC |
| HDAC6 | GCGGTGGATGGAGAAATAGA | CCGGAGGGTCCTTATCGTAG |
| HDAC8 | GCGTGATTTCCAGCACATAA | ATACTTGACCGGGGTCATCC |
| HDAC9 | GCCCACAGGAACTTCTGACT | GAACTCTAAGCCAGATGGGG |
| MGMT | GGATTGCCTCTCATTGCTCC | CCCGTTTTCCAGCAAGAGTC |
| CDKN1A | GGCAGACCAGCATGACAGATTT | AAGATGTAGAGCGGGCCTTTGA |
| PMS2 | ACTTCCGTGGATTCTGAGGG | GTGTTTGGGGTTGCGAGATT |
| RAD51 | CTTTGGCCCACAACCCATTTC | ATGGCCTTTCCTTCACCTCCAC |
| **CRISPR/Cas9** | | |
| gHDAC6 | CACCGTCCCTTGCAGTCCCACGATT | AAACAATCGTGGGACTGCAAGGGAC |
| gMGMT | CACCGTCTGCACGAAATAAAGCTCC | AACGGAGCTTTATTTCGTGCAGAC |

**Table S4.** List of antibodies used in the study with their respective dilutions.

| Antibody target | Brand/Cat. No. | Dilution |
|---|---|---|
| HDAC6 | Cell Signaling/7558 | 1:1000 |
| ac-α-tubulin | Santa Cruz/sc-23950 | 1:5000 |
| β-actin | Abcam/8227 | 1:10000 |
| H3K9ac | Cell Signaling/9649 | 1:1000 |
| H3K14ac | Cell Signaling/7627 | 1:1000 |
| H3K18ac | Cell Signaling/13998 | 1:1000 |
| H3K27ac | Cell Signaling/8174 | 1:1000 |
| H3 total | Abcam/ab9049 | 1:1000 - 1:5000 |
| cleaved PARP | Cell Signaling/9541 | 1:1000 |
| GAPDH | Abcam/ab9485 | 1:5000 |
| ATM | Cell Signaling/2873 | 1:1000 |
| p-ATM | Cell Signaling/13050 | 1:1000 |
| ATR | Cell Signaling/2790 | 1:1000 |
| p-ATR | Cell Signaling/2853 | 1:1000 |
| p21 | Cell Signaling/2947 | 1:1000 |
| Lamin B1 | Abcam/ab16048 | 1:5000 |
| MGMT | Cell Signaling/2739 | 1:1000 |
| Rabbit secondary (HRP) | Abcam/ab97051 | 1:5000 |
| Mouse secondary (HRP) | Abcam/ab97023 | 1:5000 |
| IRDye 800CW anti-rabbit | LICOR/926-32211 | 1:5000 |
| IRDye 680RD anti-mouse | LICOR/926-68070 | 1:5000 |

.
